# A bioluminescence-based chemical screen identifies a bactericidal naphthalene scaffold targeting MmpL3 in *Mycobacterium abscessus*

**DOI:** 10.1101/2025.10.26.684512

**Authors:** Samsher Singh, Ria Sorayah, Yushu Chen, Claire V. Mulholland, Wassim Daher, Carmen J. E. Pee, Tan Ai Zhu Casandra, Denise Wee, Stefan H. Oehlers, Kimberly A. Kline, Shu Sin Chng, Michael Berney, Laurent Kremer, Garrett Moraski, Kevin Pethe

## Abstract

*Mycobacterium abscessus* pulmonary disease (Mabs-PD) presents a significant and growing global health threat, particularly in individuals with underlying lung conditions like cystic fibrosis and chronic obstructive pulmonary disease. A key challenge in treating Mabs-PD is the lack of bactericidal antibiotics effective at therapeutically relevant concentrations, underscoring an urgent need for drug discovery. Targeting cell-wall synthesis is a promising approach, as evidenced by the success of broad-spectrum β-lactam antibiotics and the frontline antituberculosis drug isoniazid. However, these agents exhibit limited efficacy against Mabs, often requiring concentrations unachievable in lung tissues. Here, we used a bioluminescence-based whole-cell assay optimized to identify drugs targeting both cell-wall synthesis and the oxidative phosphorylation pathway. Screening a small drug library against Mabs revealed multiple hits, including β-lactam antibiotics, validating the effectiveness of this approach to identify cell wall-targeting agents. Among these, we identified a chemically tractable naphthalene scaffold with potent bactericidal activity. The optimized derivative GM47-1 targets MmpL3, disrupting cell wall integrity, inducing ATP leakage into the extracellular milieu, and uncoupling respiration, predominantly through the cytochrome *bcc:aa_3_* branch. Further chemical optimization resulted in a new derivative exhibiting a nanomolar minimum inhibitory concentration, with potent activity against intracellular Mabs and in a zebrafish model of infection. This study offers a promising scaffold for future therapeutic development and highlights the utility of this approach as a rapid assay platform for identifying bactericidal compounds against Mabs.

## INTRODUCTION

The rising incidence of infections caused by nontuberculous mycobacteria (NTMs) is a significant global health concern, particularly in developed countries, where these infections are becoming increasingly prevalent^1^. NTMs are environmental organisms, commonly found in soil and water, that can cause opportunistic infections in patients with underlying conditions such as chronic obstructive pulmonary disease (COPD) or cystic fibrosis (CF). *Mycobacterium avium* and *Mycobacterium abscessus* (Mabs) complexes are associated with most NTM infections in humans. *M. abscessus* pulmonary diseases (Mabs-PD) are tough to treat due to an extensive arsenal of intrinsic and acquired drug resistance mechanisms against a broad range of antibiotics and antituberculosis drugs^2–4^. The lack of approved drug regimens for Mabs-PD explains the need for lengthy chemotherapy, which may span from 6 to 24 months, associated with a low treatment success rate of 30-50%^5,6^. The recommended treatment regimen usually includes the use of a macrolide (clarithromycin or azithromycin), an aminoglycoside (amikacin), and an injectable β-lactam (cefoxitin or imipenem).

The lack of bactericidal antibiotics capable of effectively killing Mabs at therapeutically relevant concentrations^7^ is a significant limitation in the current treatment of Mabs-PD. This highlights the urgent need to discover bactericidal drugs and/or new drug regimens for treating these infections. A promising target area for the development of such drugs is cell-wall synthesis. For example, broad spectrum β-lactam antibiotics exert their bactericidal effect by inhibiting peptidoglycan synthesis, while the frontline antituberculosis agent isoniazid displays rapid bactericidal potency against *M. tuberculosis* both *in vitro* and in humans. However, antituberculosis drugs targeting cell-wall synthesis are largely ineffective against Mabs. Although some β-lactam antibiotics exhibit potency against Mabs, they require concentrations exceeding 50 mM to be effective *in vitro*^8–10^, which is not achievable in lung tissues, or need to be combined with a β-lactamase inhibitor^11^.

To accelerate the discovery of bactericidal drugs targeting cell-wall synthesis in Mabs, a rapid and effective screening assay is needed. Recently, we and others have demonstrated that drugs targeting cell-wall synthesis trigger an ATP burst in mycobacteria, easily detected using a luciferase-based microbial cell viability assay^12–14^. Although this bioluminescence signal was initially interpreted as a biological phenomenon, we demonstrate in the companion paper (see Mulholland et al., 2025, bioRxiv 2025.10.24.684413) that it results from enhanced lysis upon cell wall disruption, rather than a physiological increase in ATP levels. This mechanistic clarification enabled the development of a bioluminescence-based whole-cell assay with a dual readout, facilitating the identification of both cell wall- and bioenergetics-targeting agents in mycobacteria. The latter represents a highly relevant target space in antimycobacterial drug discovery^12,15–18^.

In this study, we adapted this whole-bacteria bioluminescence assay to identify drugs that target cell-wall synthesis (*i.e.*, those that increase the bioluminescence signal) and the oxidative phosphorylation pathway (*i.e.*, those that decrease the bioluminescence signal) in Mabs. This assay was validated and used as a dual-purpose screen against a small library of pharmacologically active drugs, resulting in several hits that either increased or decreased the bioluminescence signal. Notably, the screen identified several β-lactam antibiotics, confirming the suitability of the assay for uncovering specific cell wall-targeting agents. Among these hits, we focused on a novel chemical scaffold that induced bioluminescence deregulation associated with concomitant growth inhibition and bactericidal activity. Initial structure-activity relationship (SAR) studies demonstrated that this series is chemically tractable, leading to the identification of GM47-1, which showed improved potency compared to the initial hit and exhibited bactericidal activity both *in vitro* and against intracellular Mabs. Mode of action studies revealed that GM47-1 targets the mycolic acid transporter MmpL3, a validated drug target in both *Mycobacterium tuberculosis*^19^ and Mabs^20,21^. Interestingly, we demonstrated that the increase in bioluminescence signal was an artefact due to cell-wall weakening, which induced ATP release into the extracellular milieu. Furthermore, we showed that GM47-1 also acts as an uncoupler, leading to a decrease in intracellular ATP levels and a cytochrome *bcc:aa_3_*-dependent increase in oxygen consumption. Further chemical optimization led to the identification of a derivative with enhanced potency against intracellular Mabs.

## RESULTS

### Application of a dual-purpose whole-cell assay to identify inhibitors targeting the cell-wall and energy metabolism

In a quest to identify novel chemical series for Mabs-PD, we used a simple ATP bioluminescence-based method developed for tuberculous and non-tuberculous mycobacteria (Mulholland et al., bioRxiv 2025.10.24.684413), to identify drugs targeting either cell-wall synthesis (*i.e.*, drugs triggering an increase in bioluminescence^13,14,22^) and energy metabolism inhibitors (*i.e.*, drugs triggering a reduction in bioluminescence^15,22,23^). The screening was performed in 96-well plate format against MabsΔ*cydABDC* using the BacTiter-Glo^TM^ luminescence-based assay kit. Mabs, similar to *M. tuberculosis*, possesses two terminal oxidases: the cytochrome *bcc:aa₃* (cyt-*bcc:aa_3_*) and the cytochrome *bd* oxidase (cyt-*bd*)^15,24^. The rational for using the MabsΔ*cydABDC* deficient for the expression of the cytochrome *bd* oxidase, was to favour the identification of drugs specifically targeting cyt-*bcc:aa_3_*, a clinically validated target for tuberculosis^18,25–27^. For assay development, bedaquiline (BDQ) was used as a positive control to inhibit ATP production, while cefoxitin (β-lactam) was included as a positive control to induce an increase in ATP bioluminescence^13,23^. The z’ score for ATP depletion and ATP signal increase were 0.62 and 0.91, respectively using BDQ and cefoxitin (Figure 1A & 1B), indicating the robustness of the assay to identify relevant hits. A chemical screen was conducted using the Sigma Library of Pharmacologically Active Compounds (LOPAC**^®1280^**), comprising 1,280 known drugs. A total of seven drugs decreased ATP levels by at least 50% relative to basal ATP levels in untreated samples (Figure 1C; Table S1). In a dose response assay, five of the seven hits demonstrated ATP IC_50_ in the range of 2.5-25.9 µM against MabsΔ*cydABDC*. When tested against the parental Mabs strain, all ATP-depleting hits showed comparable IC_50_ values, suggesting that their activity was not dependent on the absence of cyt-*bd* oxidase. This lack of selectivity for the Δ*cydABDC* strain indicated that these compounds did not preferentially target the cyt-*bcc:aa_3_*, and they were therefore ruled out from subsequent studies. A total of eleven drugs induced an increase in bioluminescence of at least 50% relative to the DMSO (vehicle) control (Figure 1; Table 1). Two drugs that were ATP analogues (2-chloroadenosine triphosphate and P^1^,P^4^-Di(adenosine-5’) tetraphosphate) were discarded due to assay interference and are not listed in the table. Validation in the Mabs parental strain demonstrated a comparable increase in bioluminescence (Table S2). The validated hit list contained an over-representation of antibiotics targeting peptidoglycan synthesis, supporting the assay’s utility in identifying cell wall-targeting agents. The nine hits induced a bioluminescence increase ranging from 1.6 to 4.8-fold, except for SBI-0087702, which triggered a much higher signal increase up to 52.7-fold relative to the untreated control (Table 1). Next, the hits were confirmed in a dose-response assay for both bioluminescence increase and growth inhibition. The ATP effective concentration 50% (ATP EC_50_) ranged from 1.0 to 22.9 µM against MabsΔ*cydABDC* and had a comparable effect on the luminescence signal against the parental strain (Table S2). Despite the significant increase in bioluminescence signal, none of the peptidoglycan-targeting antibiotics were growth inhibitory up to a concentration of 40 µM. This result may indicate that these drugs inflicted limited cell wall damage that was sufficient to induce a bioluminescence increase, but not to inhibit bacterial growth. The only hit that was growth inhibitory was SBI-0087702 (Table S2). In-house resynthesized SBI-0087702 (renamed GM46-96; Fig 2A) demonstrated a potency comparable to the hit compound and exhibited an inverse correlation between bioluminescence increase and growth inhibition (Figure 2B).

**Figure 1.**
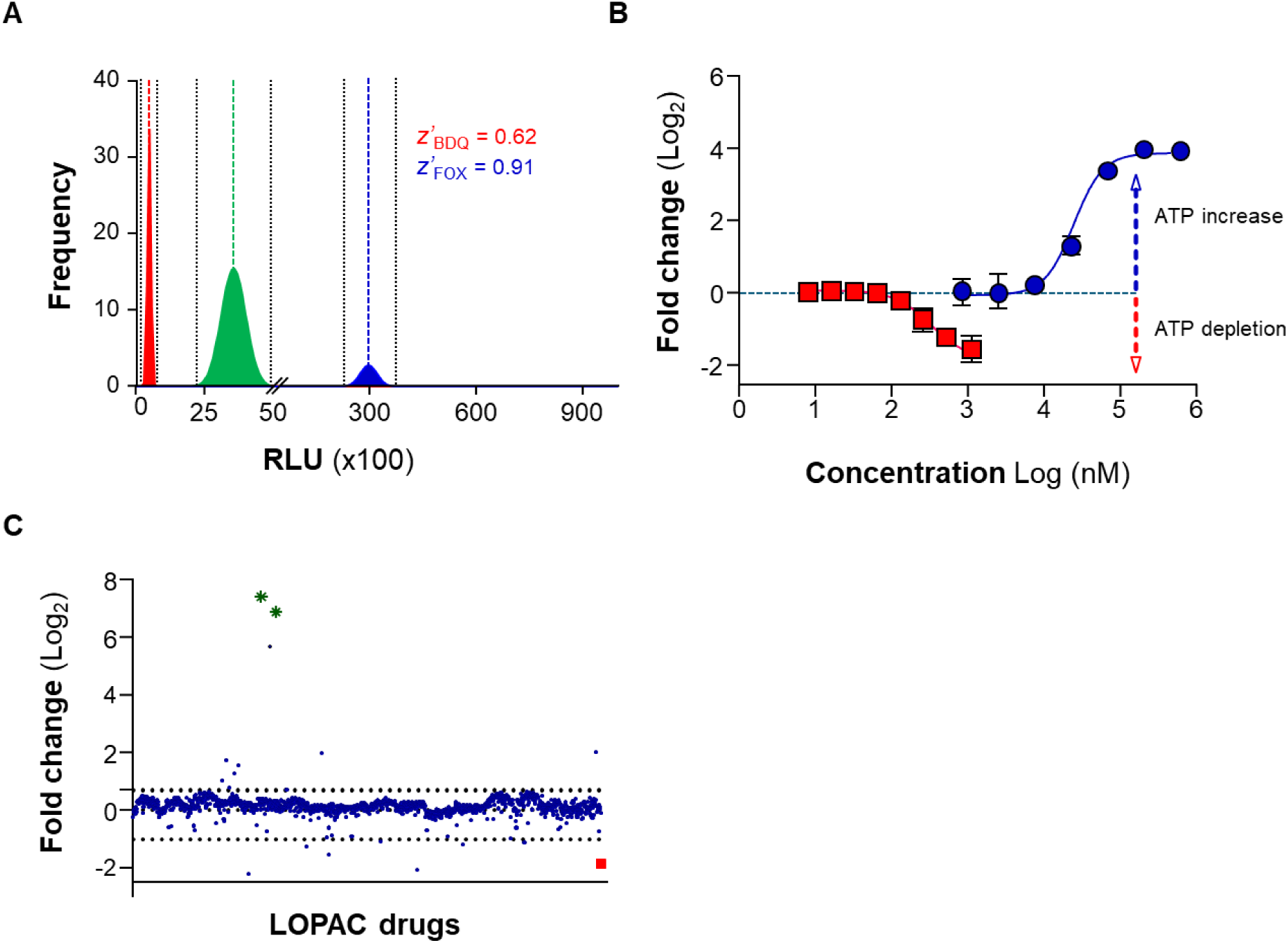
A dual ATP bioluminescence-based chemical screen against Mabs. (A) The graph shows the gaussian distribution frequency of values yielded from drug free control (green), cefoxitin (FOX; blue) and bedaquiline (BDQ; red) in the assay. Dotted lines (black) indicate standard deviations in both directions from their respective means. (B) ATP assay results showing the concentration dependent ATP increase and ATP depletion caused by cefoxitin (blue line) and BDQ (red line), respectively. (C) LOPAC**^®1280^** library was screened at a single concentration of 40 µM against Mabs Δ*cydABDC*. Bioluminescence values were expressed as fold change relative to basal ATP levels. The average value of BDQ positive control is depicted by a red square. The dotted line indicates cut-off values for selecting hits based on bioluminescence induction. The two hits shown as green stars are ATP analogues.

**Figure 2.**
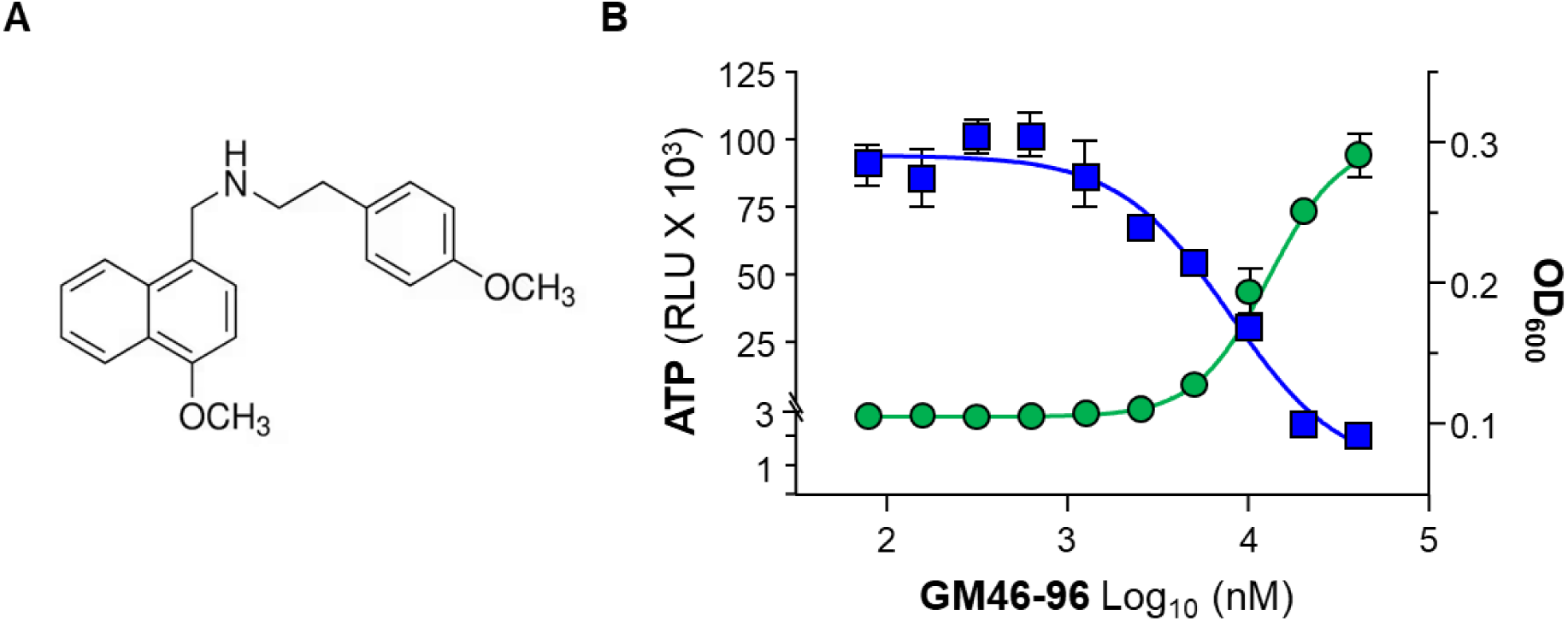
SBI-0087702 induces a bioluminescence increase and inhibits growth of Mabs. (A) Structure of SBI-0087702 (GM46-96). (B) Effect of a dose-range of GM46-96 (SBI-0087702) on bioluminescence indicative of inhibition of ATP (green circles) and growth inhibition (blue squares). For ATP and MIC determination, wild-type Mabs culture was treated with GM46-96 for 4 h and 48 h respectively.

**Table 1.** Identification of hits from LOPAC^®1280^ library screen for ATP perturbations. A total of 9 hits (drugs) were identified to trigger an ATP increase relative to basal levels against Mabs Δ*cydABDC*. One among nine hits was able to inhibit growth of Mabs. BDQ, a F_1_F_O_ ATP synthase inhibitor was used as a control drug which inhibits ATP synthesis and growth in Mabs. Basal levels: ATP basal levels in untreated controls. EC_50_ is defined as effective concentration required for half the maximal increase in bioluminescence (ATP).

| Drug name | Target | ATP<br>(× fold<br>increase) | ATP<br>EC<sub>50</sub> (μM) | MIC<sub>50</sub><br>(μM) |
| --- | --- | --- | --- | --- |
| Cephalexin | Peptidoglycan<br>(antibiotic) | 2.2 | 1.0 | >40 |
| Cefaclor |  | 3.5 | 2.1 | >40 |
| Ceftriaxone |  | 1.8 | 1.3 | >40 |
| Cefotaxime |  | 2.6 | 10.5 | >40 |
| Cephadrine |  | 3.1 | 3.0 | >40 |
| Vancomycin |  | 3.8 | 9.0 | >40 |
| SBI-0087702 | Mitochondrial ATF<br>translocator | 52.7 | 8.4 | 5.6 |
| Demeclocycline | Ribosome<br>(antibiotic) | 1.6 | 2.7 | >40 |
| NS8593 | Ca<sup>2+</sup>-activated K<sup>+</sup><br>channels | 4.8 | 22.9 | >40 |

### GM46-96 is amenable to chemical optimization

Given the modest growth inhibitory potency of GM46-96, twelve naphthalenemethanamine derivatives were synthesized to evaluate chemical tractability. The derivatives were evaluated for their effects on bioluminescence increase and growth inhibition. Five of the twelve derivatives were potent and displayed a good correlation between ATP EC_50_ and MIC_50_ (Figure 3A and 3B). The SAR around the naphthalenemethanamine hit (SBI-0087702 or GM46-96; Fig 2A) revealed that the ethyl linkage is optimal for potency, as the benzyl (MSU-06) and aniline (MSU-21) derivatives were both inactive (MIC_50_ >20 µM). Even the 4-(4-methoxyphenoxy)aniline (MSU-19) was inactive, suggesting that the amine “sidechain” appears to be limited in size to accommodate the binding pocket. The optimal size chain hypothesis seems to be corroborated by the weak activity observed in GM46-72, which bears the highly dynamic and rotatable 3-phenylpropan-1-amine moiety (MIC_50_ = 30 μM) but is one carbon longer. It was also found that neither the amide (MSU-43) nor the directly coupled amine (MSU-134) is tolerated (MIC_50_ > 40 µM, respectively) in this binding pocket. The MIC improved when the 4-methoxy was substituted by the 4-ethoxy moiety, giving rise to GM47-1 with an MIC_50_ of 1.6 µM. The placement of this enhanced substituent was evaluated by preparing the 3-ethoxy (MSU-24) and 2-ethoxy (MSU-26) derivatives, which revealed that potency was reduced with the 3-ethoxy (MSU-24, MIC_50_ = 2.3 μM) and abolished with the 2-ethoxy (MSU-026, MIC_50_ > 20 μM). Also found detrimental was the replacement of the methoxy group on the naphthalene core with a hydroxyl group (as in MSU-40, MIC_50_ > 40 μM). Exploration of alternative substituents upon the side chain resulted in the identification of two active analogs. The 4-chloro analog (GM46-75) had an MIC_50_ = 3.3 μM, while 4-trifluoromethoxy analog (MSU-18) had an MIC_50_ = 2.5 μM. While neither was superior to GM47-1 (bearing the 4-ethoxy moiety), they demonstrate that the choice of substituent does impact potency.

**Figure 3.**
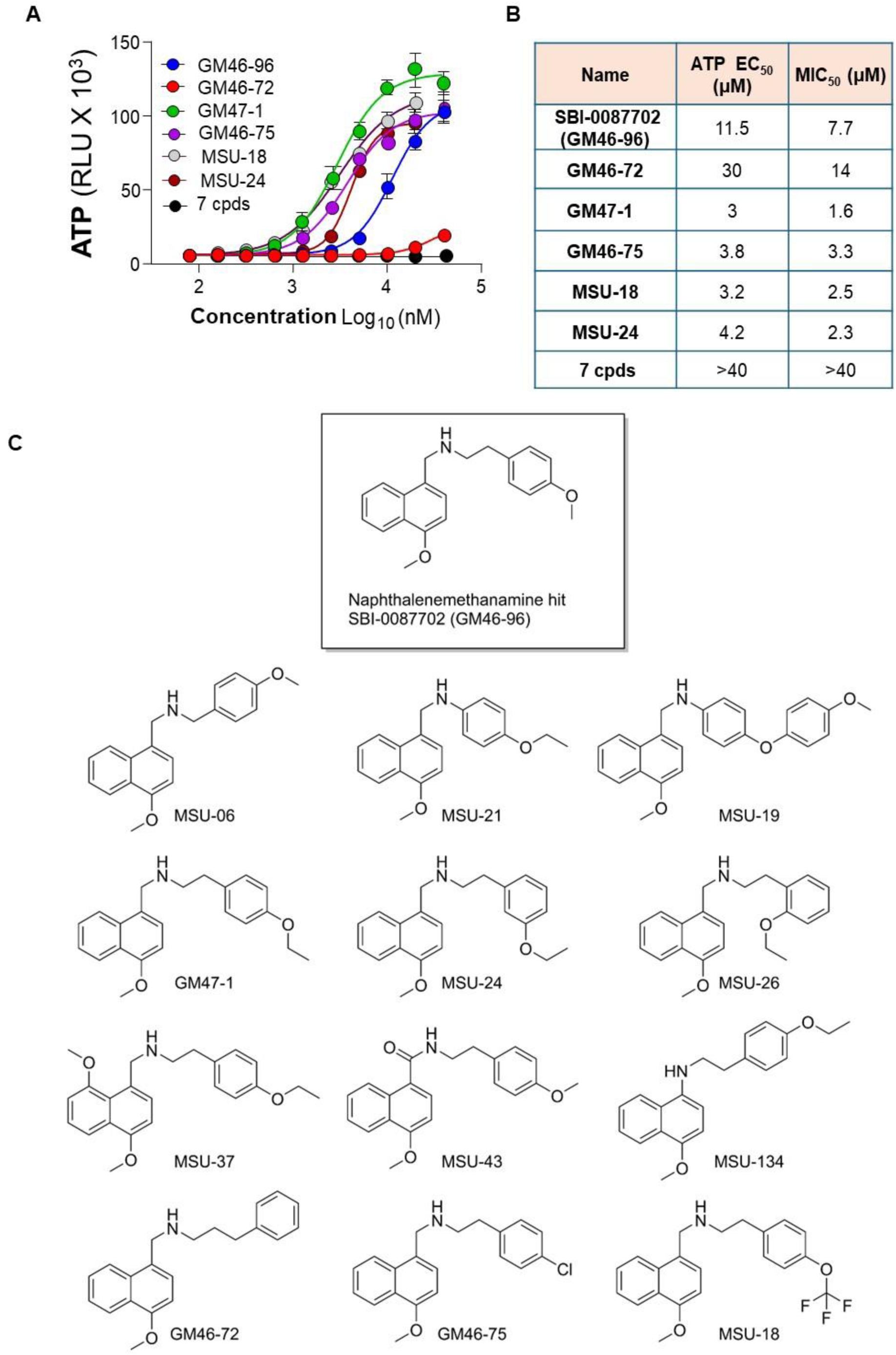
SBI-0087702 is amenable to chemical optimization. (A&B) GM46-96 and 12 chemically synthesized derivatives were assessed for ATP deregulation against Mabs. Active hits trigger a dose-dependent ATP increase that was inversely correlated with growth inhibition. (C) SAR around naphthalenemethanamine hit, SBI-0087702, to arrive at most potent compounds GM47-1, GM46-75, MSU-18, and MSU-24.

### GM47-1 is potent against clinical isolates, bactericidal *in vitro* and potent against intracellular Mabs

A critical factor in antibacterial development relies on the potency of the tested compound against clinical isolates, including those resistant to clinically used antibiotics. GM47-1 was assessed against a panel of Mabs clinical isolates, using BDQ as a reference drug. All clinical isolates tested were susceptible to GM47-1, regardless of their antibiotic-resistance profile (Table S3), indicating a mode of action distinct from approved antibiotics. Since the objective of the screening effort was to identify bactericidal drugs, we conducted kill kinetics experiments. Mabs CIP104536^T^ was exposed to GM47-1 at various concentrations, and CFUs were enumerated over time. BDQ was used as a control bacteriostatic agent. The results revealed that GM47-1 at 6 ×MIC_50_ killed around 99.2% and 99.8% of the initial inoculum in 2 and 5 days, respectively, demonstrating that the drug is rapidly bactericidal *in vitro* (Figure 4A). Like other pathogenic mycobacteria, Mabs invade phagocytic cells and persist intracellularly, a property that may aid immune and antibiotic evasion^28^. For instance, intracellular mycobacteria are protected mainly from amikacin, an antibiotic that penetrates poorly in eukaryotic cells^29^. The ability of GM47-1 to target intracellular Mabs was evaluated in THP-1 macrophages. Both clinically used antibiotics BDQ and clarithromycin were bacteriostatic in this model (Figure 4B). GM47-1 was potent against intracellular Mabs, reducing bacterial count in a dose-dependent manner and statistically more efficiently than BDQ (Figure 4B). This result indicates that GM47-1 can penetrate eukaryotic cells and that its target is essential for the viability of intracellular Mabs.

**Figure 4.**
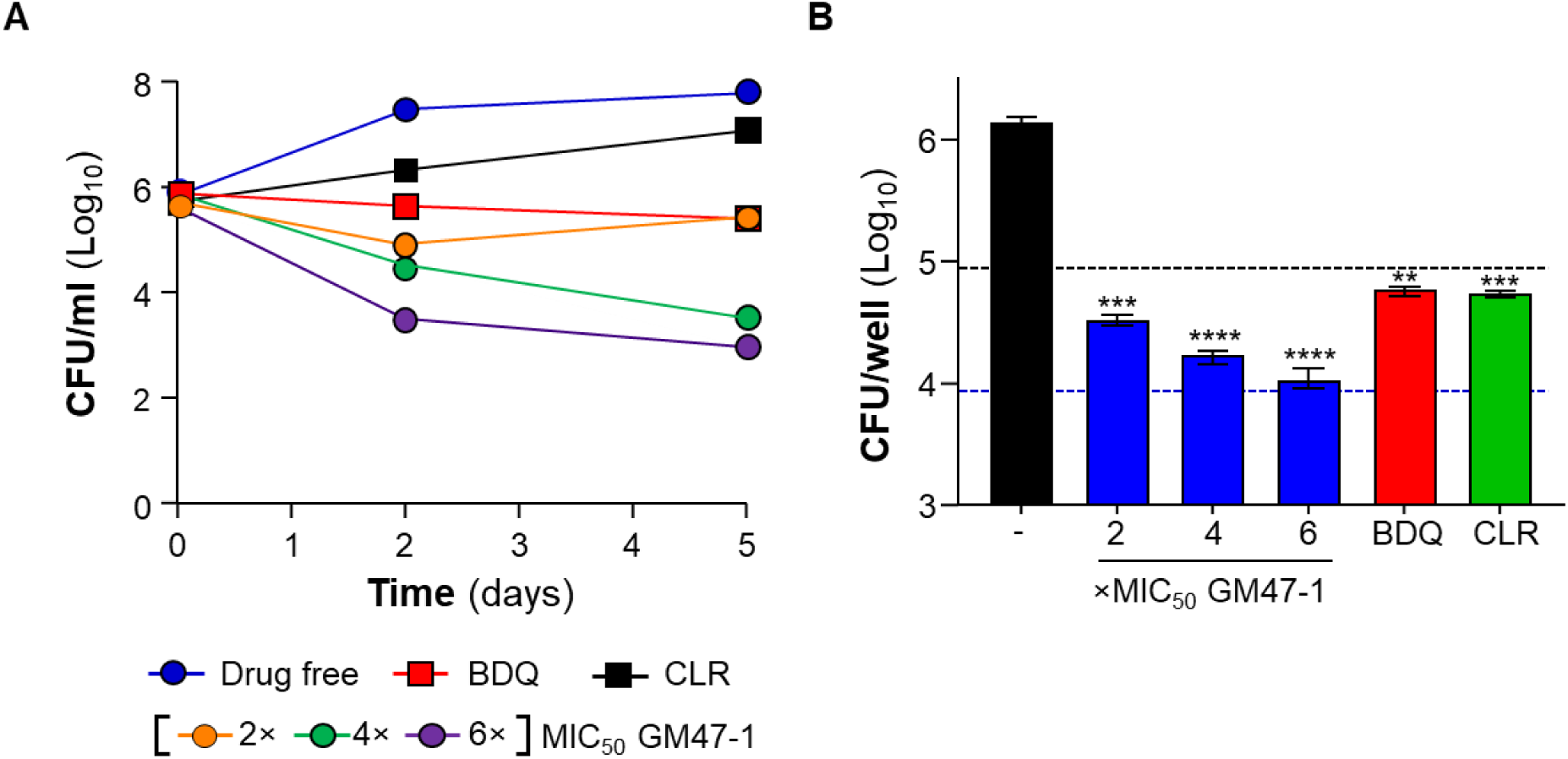
GM47-1 is bactericidal and potent against intracellular Mabs. (A) Kill kinetics of GM47-1 at 2-6 ×MIC_50_. BDQ (MIC_50_ of 0.2 µM) and clarithromycin (MIC_50_ of 0.5 µM) were used as drug control. Bacterial viability was determined by enumerating colony-forming units (CFU) after two- and five-days post-treatment. Drug free indicates the vehicle only control *i.e.* 1% DMSO. (B). Differentiated THP-1 cells were infected with M. abscessus at an MOI of 1. At an interval of 1 h post-infection, cells were treated with 2-6 ×MIC_50_ of GM47-1, 1% DMSO (vehicle control; black), BDQ at 10 ×MIC_50_ (2 µM), or clarithromycin (CLR) at 20 ×MIC_50_ (10 µM). Cells were treated for 48 h before lysis for bacterial viability assessment by CFU counts on agar plates. Upper and lower dotted lines indicate input cells at the start of treatment and 90% reduction in cells respectively. The data points are expressed as mean ± SD of triplicates of a representative experiment. \**P* < 0.05; \*\**P* < 0.01 unpaired Student’s t-test, two-tailed; n = 3; comparing inoculum *vs* treatment groups. Exact *P*-values (from left to right): 0.0001, <0.0001, <0.0001, 0.0017 and 0.0009.

### GM47-1 targets MmpL3 in Mabs

To understand the mechanism of action of GM47-1, we analyzed the profiles of [^14^C]-labelled newly synthesized lipids extracted from Mabs cells upon drug treatment for 1.5 h. GM47-1 up to ∼5x MIC (16 µM) did not affect *de novo* mycolic acid biosynthesis in the cells (Figure S1). However, GM47-1 treatment resulted in markedly lower production of trehalose dimycolates (TDM), with a concomitant accumulation of trehalose monomycolates (TMM). Furthermore, the drug also inhibited the attachment of mycolates to arabinogalactan-peptidoglycan (Figure 5A), suggesting that GM47-1 abolishes TMMs transport to the outer membrane, consistent with possible inhibition of the trehalose monomycolate transporter MmpL3^20,30,31^.

**Figure 5.**
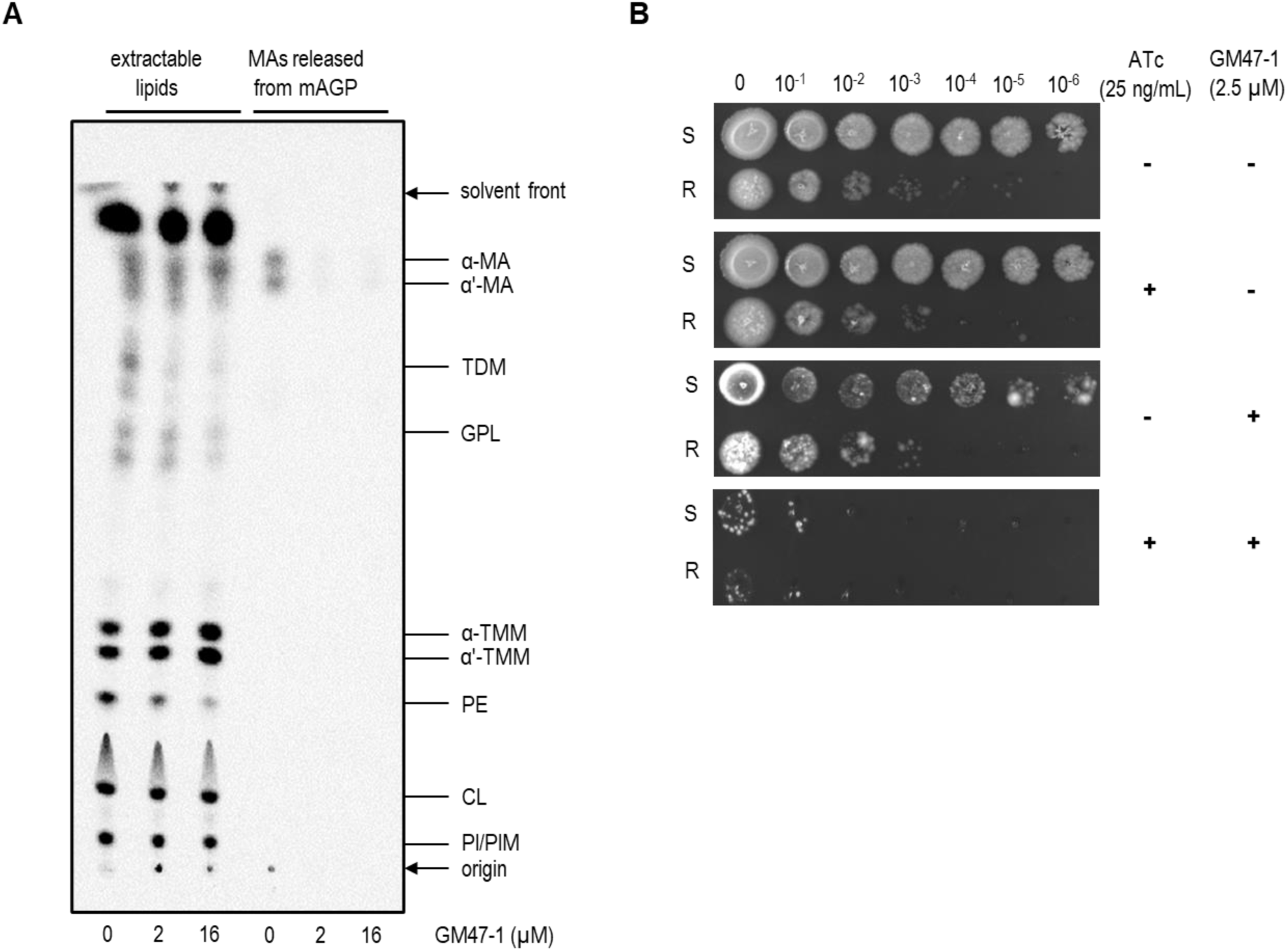
GM47-1 blocks TMM transport to the outer membrane and Mabs growth susceptibility is *mmpL3*-dependent. (A) TLC analysis of [^14^C]-labeled total extractable lipids (left three lanes) and MAs released from mAGP (right three lanes) extracted from Mabs cells treated with indicated concentrations of GM47-1. The same amount of radioactivity was loaded for each extractable lipid sample, while released MAs were loaded in the same volume as extractable lipids from the same cells. The TLC was visualized by phosphor imaging. Each TLC is representative of at least two biological replicates. GPL, glycopeptidolipid; CL, cardiolipin; PE, phosphatidylethanolamine; PI, phosphatidylinositol; PIM, phosphatidylinositol mannoside; MA: mycolic acid; TDM: trehalose dimycolate; TMM: trehalose monomycolate. (B) Smooth (S) and rough (R) *mmpL3* conditional mutants were grown to exponential phase, and 3 µL of 10-fold serial dilutions were spotted onto LB agar medium supplemented or not with 25 ng/mL ATc in the presence or not of 2.5 µM GM47-1. Plates were incubated at 37 °C and pictures taken after 3 days.

To validate the molecular target of GM47-1, we selected escape mutants resistant to the compound on agar plates. The estimated spontaneous rate of mutations to GM47-1 was 1.6×10^−7^ and 2.6×10^−8^ when tested at 1.2-fold and 2.3-fold the agar MIC, respectively. These results indicated a low probability of emergence of resistant mutants compared with approved antimycobacterial drugs^32^ and as reviewed for diverse MmpL3 inhibitors^33^. Fourteen colonies were picked for further characterization. Upon confirmation of stable resistance to GM47-1 that ranged from 3.6 to >19-fold higher MICs compared to the parental strain, all colonies were subjected to whole-genome sequencing. Single-nucleotide substitutions leading to single amino acid changes (Table 2) were identified in the *mmpL3*-encoding gene *MAB_4508* in all colonies. MmpL3 is a validated drug target in *M. tuberculosis*^19^ and Mabs^20,21^, and is required for shuttling trehalose monomycolate (TMM) across the mycobacterial plasma membrane, where it is utilized by the Ag85 complex to mycolylate arabinogalactan (AG) or to produce trehalose dimycolate (TDM)^31,34,35^. Among the five substitutions that were identified, the three most common (L260P, V299A, and A309P) are involved in resistance to other classes of MmpL3 inhibitors^20,30,31^. To the best of our knowledge, the substitutions A662V and L692F have not yet been implicated in resistance to MmpL3 inhibitors.

**Table 2.** Characteristics of GM47-1 spontaneous resistant mutants against Mabs. MIC_50_ was determined against a panel of resistant mutants. Whole genome sequencing led to identification of single nucleotide polymorphisms (SNPs) in *MAB_4508* and corresponding amino acid changes (mutation harboring transmembrane helices) are also indicated.

| Strains # | GM47-1<br>(MIC<sub>50</sub>, µM) | BDQ<br>(MIC<sub>50</sub>, µM) | MAB_4508<br>DNA SNPs | AA change<br>(TM helices) |
| --- | --- | --- | --- | --- |
| Parental | 2.1 | 0.31 | - | - |
| 1 | >40 | 0.24 | T779C | L260P (TM-4)<sup>a</sup> |
| 2 | 18.0 | 0.19 |  |  |
| 3 | >40 | 0.59 |  |  |
| 4 | >40 | 0.27 |  |  |
| 5 | >40 | 0.27 |  |  |
| 6 | 11.0 | 0.10 | T896C | V299P (TM-5)<sup>b</sup> |
| 7 | >40 | 0.30 | G925C | A309P (TM-5)<sup>a,c</sup> |
| 8 | >40 | 0.25 |  |  |
| 9 | >40 | 0.25 |  |  |
| 10 | 26.4 | 0.25 |  |  |
| 11 | >40 | 0.3 |  |  |
| 12 | >40 | 0.25 |  |  |
| 13 | 15.4 | 0.21 | C1985T | **A662V** (TM-11)<sup>d</sup> |
| 14 | 7.6 | 0.37 | C2074T | **L692F** (TM-12)<sup>d</sup> |

<sup>a</sup> Dupont et al., 2016; Mol Microbiol. <sup>20</sup>. <sup>b</sup> Raynaud et al., 2020; ACS Infectious Disease <sup>30</sup>. <sup>c</sup> Kozikowski et al., J. Med. Chem<sup>31</sup>, <sup>d</sup> highlighted in bold: novel mutations.

To further confirm that MmpL3 is the primary target of GM47-1, we employed a Mabs *MAB_4508* hypomorph strain^36^. This conditional knock-down mutant, based on a Tet-Off system, was previously engineered to demonstrate the essentiality of *mmpL3* for Mabs growth both *in vitro* and in a zebrafish model of infection^36^. In this system, *mmpL3* expression is repressed in an anhydrotetracycline (ATc)-dependent manner. While high concentrations of ATc markedly inhibit growth on agar, a lower concentration of 25 nM ATc has minimal effect on its own. We therefore used this sublethal ATc concentration to assess the sensitivity of the knock-down strain to GM47-1. Supplementation with 2.5 µM GM47-1 caused only mild growth inhibition in the absence of ATc (Figure 5B). In contrast, when combined with 25 nM ATc, GM47-1 almost completely abolished growth of the *mmpL3* hypomorph in both smooth (S) and rough (R) morphotypes (Figure 5B). These findings indicate that partial repression of *mmpL3* expression sensitizes Mabs to GM47-1, providing further evidence that MmpL3 as its likely target.

### GM47-1 induces extracellular ATP release and uncouples respiration

GM47-1 stood out in the primary screen by increasing the bioluminescence signal about 50-fold compared to untreated controls (Figure 3A). This, together with its elucidated target, MmpL3, indicated that GM47-1 weakens the Mabs cell wall. We therefore sought to validate the basis for the increased ATP signal and its connection to the bactericidal potency of the drug. To begin, we exploited a Mabs strain expressing an ATP biosensor (Mab mDB158)^37^ to measure intracellular ATP levels in response to GM47-1 treatment. In sharp contrast to the BacTiter-Glo^TM^ assay results, GM47-1 reduced the intracellular ATP pool in the reporter strain in a dose-dependent manner (Figure 6A). This further independently confirmed that the observed ATP-burst detected by BacTiter-Glo^TM^ results from enhanced ATP release rather than an actual increase in intracellular ATP levels (Mulholland et al., bioRxiv 2025.10.24.684413). We next measured extracellular ATP to determine if GM47-1 lyses the cells directly, or, if the ATP release is the result of synergism between cell wall weakening and lysis buffer action. Results showed that the culture broth medium of GM47-1-treated bacteria contained more ATP than untreated control, suggesting that the drug sufficiently disrupts the cell envelope to allow for ATP to leak out, a phenomenon not observed with amikacin (Figure 6B). Inhibition of ATP synthesis upon binding to MmpL3 could be explained by oxidative phosphorylation uncoupling. A hallmark of uncoupling is the stimulation of the electron transport chain to generate a protonmotive force, which is subsequently dissipated by the uncoupling agent. We tested this hypothesis by measuring the oxygen consumption rate (OCR) of the bacterium upon drug treatment. Using a semi-quantitative methylene blue assay^15^, we observed that GM47-1-treated cultures decolorized within 24 h, whereas the untreated control remained blue at 48h (Figure 6C), and required 72 h to fully decolorize (Figure S2). As a control, we confirmed that BDQ inhibited oxygen consumption in Mabs (Figure 6C), consistent with previous findings in *M. tuberculosis*^15^. Next, we measured the OCR using the Seahorse XFe96 platform. In this assay, GM47-1 kinetically increased OCR (Figure 6D) by 1.48-fold at its MIC concentration in 2 h. While BDQ also increased OCR in this assay (Figure 6D), as previously reported in *M. tuberculosis*^38^, this effect appeared to be transient, with the drug ultimately inhibiting OCR in the long term (Figure 6C). Mabs expresses two aerobic respiratory branches: the cyt-*bcc:aa_3_* (encoded by *qcrCAB*) and the cyt-*bd* oxidase (*cydABDC*). To determine which branch was responsible for accelerated respiration, we made use of MabsΔ*qcrCAB* and Δ*cydABDC* strains^39^. The growth inhibitory potency of GM47-1 was unaffected by the loss of either terminal oxidase (Figure S3). While the OCR increase was unaffected in the GM47-1-treated Δ*cydABDC* strain, the drug induced only a marginal increase in OCR in the Δ*qcrCAB* strain, indicating that accelerated respiration is largely mediated by the cyt-*bcc:aa*_3_ respiratory branch (Figure S4).

**Figure 6.**
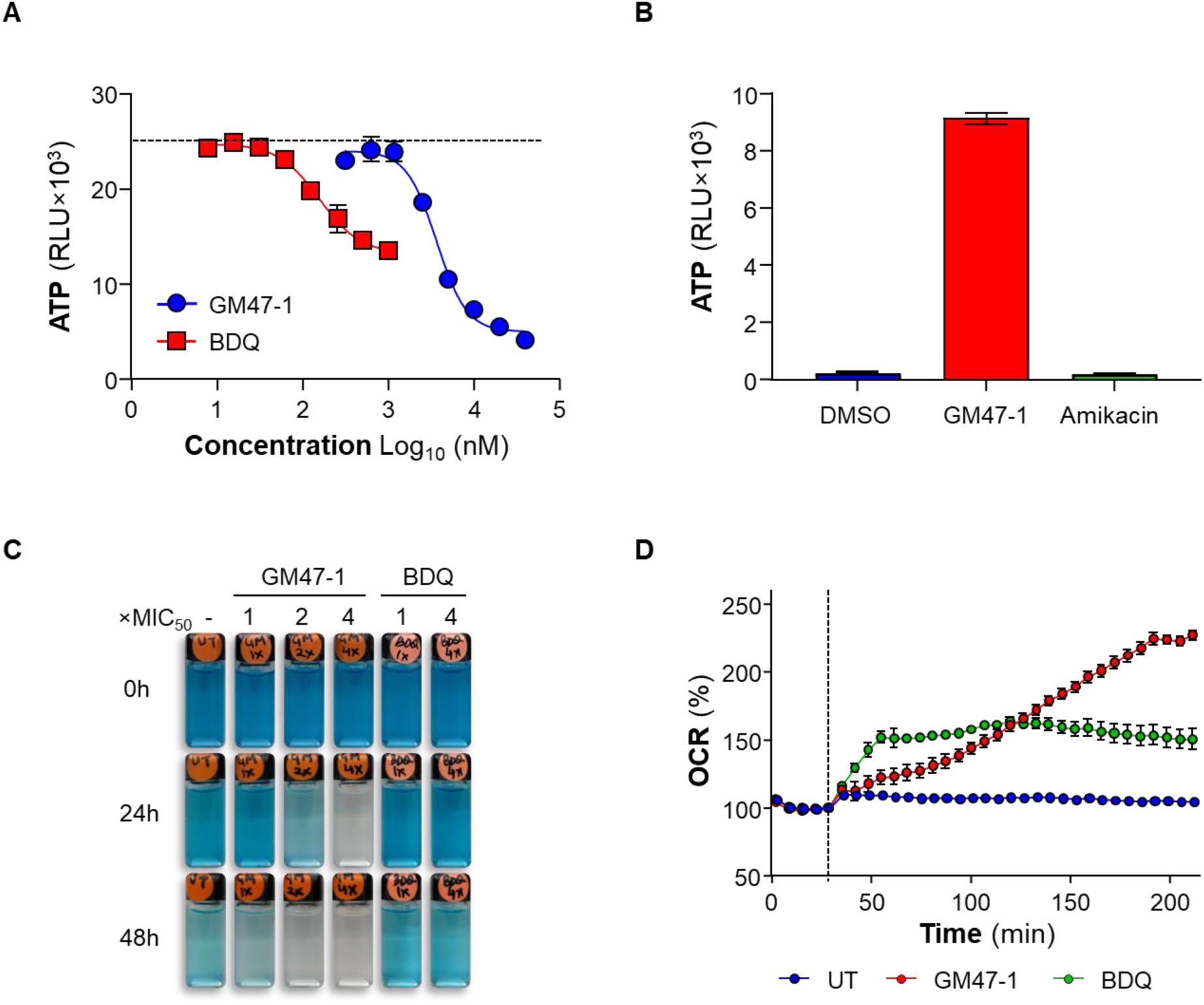
GM47-1 treatment uncouples respiration and induces ATP leakage. (A) MIC of GM47-1 and BDQ was determined against Mabs strain mDB158 expressing *lux13*. After 2 days of incubation bioluminescence was measured which is indicative of cellular ATP levels (B) Exponentially grown culture was treated with 4×MIC of GM47-1 and amikacin for 4 h, followed by measurement of ATP levels in filtered supernatant using BacTiter-Glo^TM^. (C) Qualitative measurement of oxygen consumption by Mabs using the oxygen sensor methylene blue at 0.001% (v/v). Exponentially grown culture was treated with GM47-1 at 1-4×MIC in tightly sealed glass vials. BDQ at 1× and 4× MIC served as control. (D) Oxygen consumption assay in Mabs using the Seahorse XFe96 Extracellular flux analyzer. GM47-1 and BDQ were injected at 1 ×MIC_50_. For each condition, OCR readings were normalized to the last basal OCR (fifth read) reading. OCR: oxygen consumption rate. Dotted line: injection time point.

### GM47-1 synergizes with β-lactam antibiotics

Inhibitors of MmpL3 have previously shown synergistic killing when combined with β-lactam antibiotics such as cefoxitin (FOX) and imipenem, which are used clinically to treat Mabs-PD^40^. To assess the potential of GM47-1 in combination therapy, we tested its bactericidal activity at 4 ×MIC_50_ in the presence of FOX or meropenem (MER) at growth-inhibitory concentrations. FOX alone was bactericidal, reducing the initial bacterial load, and further enhanced the killing effect of GM47-1 by 51.1% and 98.0% after 2 and 5 days of treatment, respectively (Figure 7A). MER, which lacked bactericidal activity on its own, markedly potentiated GM47-1-mediated killing by 82.4% and 98.9% at the same time points (Figure 7B). These findings are consistent with previous reports of synergistic interactions between MmpL3 inhibitors, such as indole carboxamides and β-lactams, against both *M. tuberculosis*^41^ and Mabs^40^.

**Figure 7.**
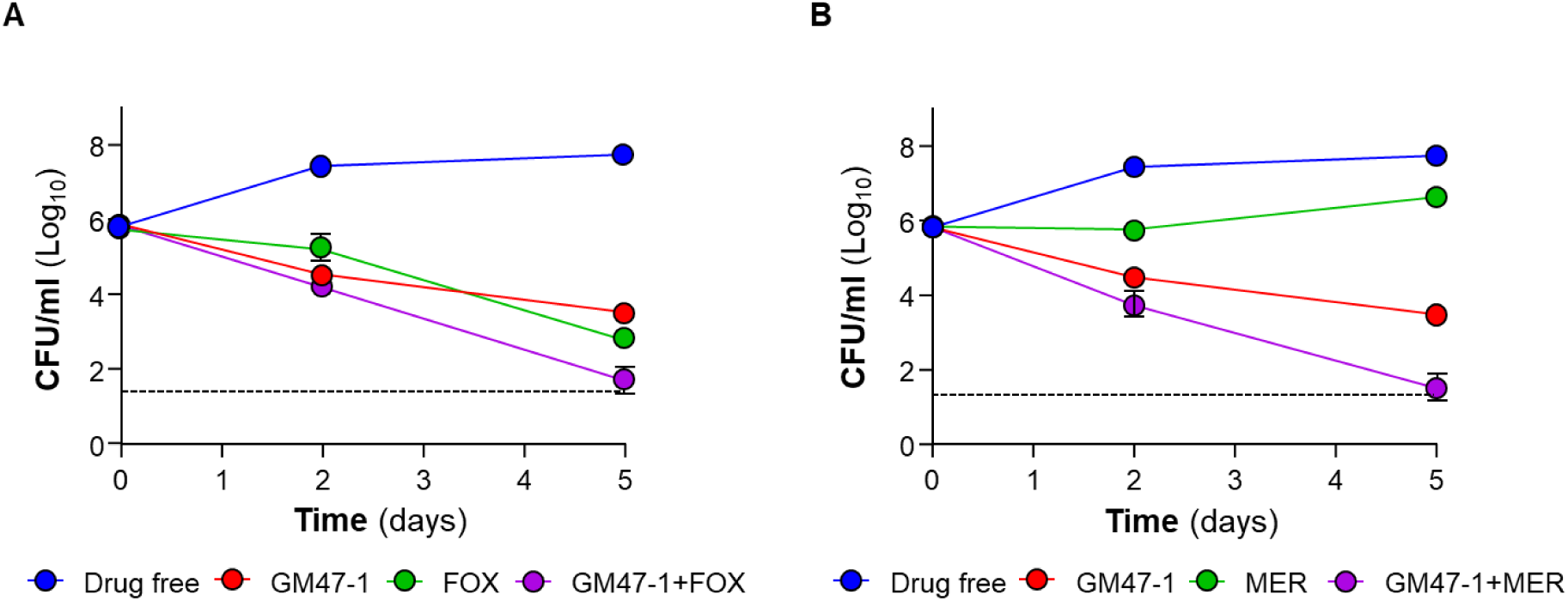
Synergistic killing of GM47-1 with cefoxitin or meropenem. Bactericidal kinetics of GM47-1 in combination with cefoxitin (FOX) or meropenem (MER) against Mabs. Bacterial viability was determined by enumerating CFU plating on agar plates. GM47-1 was used at 10 µM (4 ×MIC_50_), and FOX or MER were used at 40 and 80 µM respectively, which is equivalent to 2 ×MIC_50_ for the respective drugs. Drug free indicates the vehicle only control *i.e.* 1% DMSO. Experiments were performed in triplicate and repeated at least once. The data are expressed as the means ± SD of triplicates for each condition.

### An optimized GM47-1 derivative shows improved potency and is potent in zebrafish

New analogs were designed based on the SAR trends observed in the twelve analogs prepared and screened (Fig. 3C). The replacement of the methoxy group on the naphthalene core with a fluorine resulted in an analog, MSU-31 (Fig. 8A), with good potency (MIC_50_ = 2.05 μM). Investigating this fluorinated naphthalene core with two additional side chains revealed a significant enhancement in potency. MSU-155 (bearing a 4-trifluoromethoxy) had an MIC_50_ = 0.21 μM, and MSU-156 (bearing a 3,4-dichloro) had an MIC_50_ = 0.18 μM (Fig. 8A). Both optimized derivatives *i.e.* MSU-155 and MSU-156 exhibit a wide safety index (CC_50_/MIC_50_) of >238 and 268, respectively (Table S4). Both compounds were found to be relatively more bactericidal against intracellular Mabs when compared to GM47-1 at 4 ×MIC_50_. MSU-155 reduced the burden of intracellular Mabs in THP-1 cells by 93 and 97% and MSU-156 reduced the burden by 87 and 95% when used at 4 × and 8 ×MIC_50_, respectively (Figure 8B). Next, MSU-155 was evaluated in a zebrafish embryo-Mabs model of acute infection^42–44^. The zebrafish is a well-established model for Mabs infection that recapitulates key aspects of host–pathogen interactions and allows quantitative assessment of bacterial burden^11,23,42,43,45^. Initial tolerability experiments indicated that a concentration of 5 µM (final concentration in fish water, ∼24-fold MIC_50_) was tolerated by the fish. When treated, bacilli load as reflected by fluorescent pixel counts (FPC)^44^, showed significant reduction relative to vehicle-treated control (Fig 8C), indicating potency in this model.

**Figure 8.**
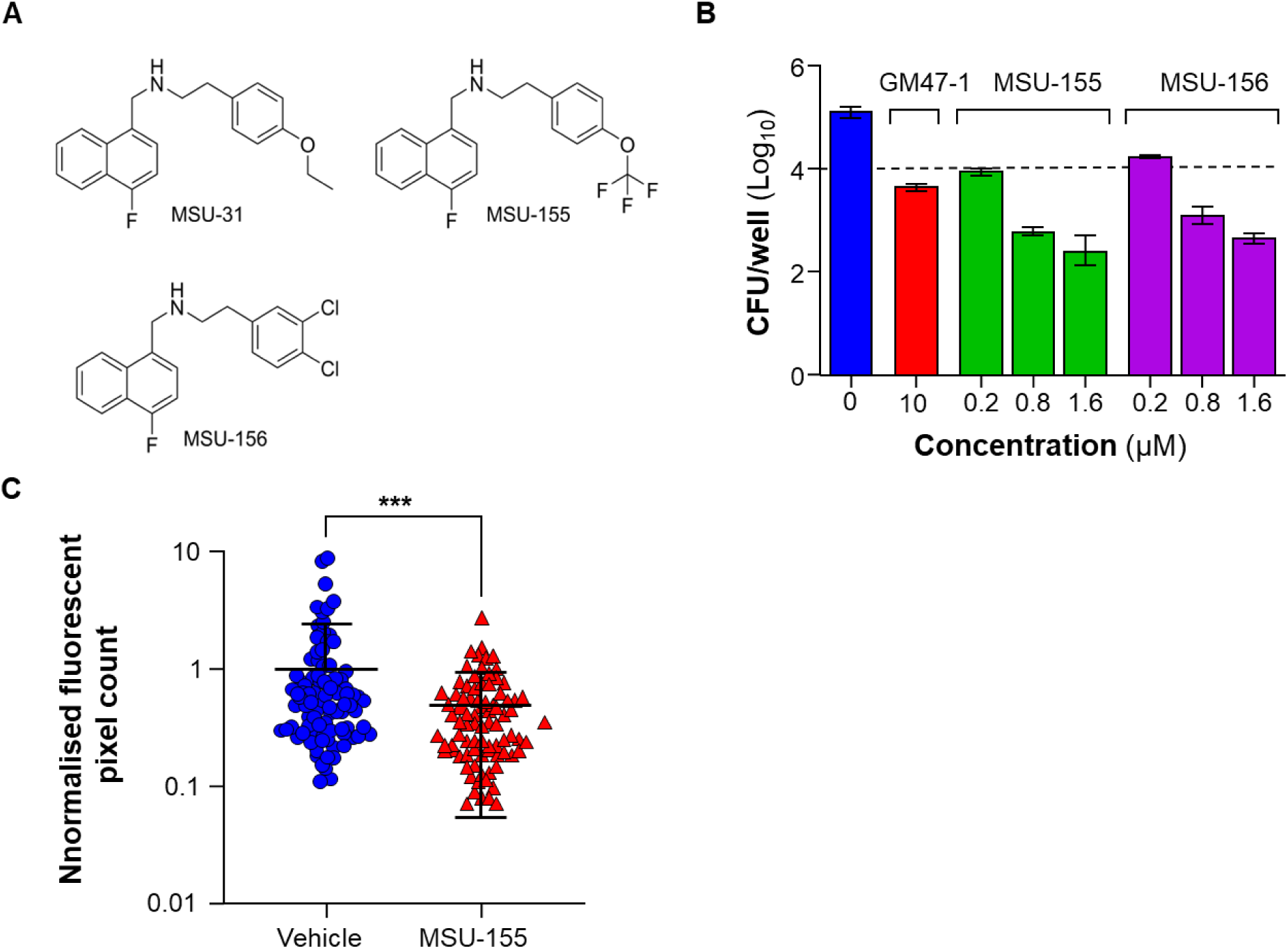
Additional SAR around GM47-1 led to enhanced potency *ex vivo*. (A) Chemical structures of GM47-1 derivatives. (B) For *ex vivo* efficacy, THP-1 macrophages were infected with Mabs. Infected macrophages were exposed to compounds for 3 days. Bacterial counts were determined by plating cell lysates onto 7H10 agar. The dotted line represents the number of bacilli internalized by macrophages before treatment. (C*) In vivo* efficacy was assessed using infecting zebrafish embryos by microinjection, followed by treatment using MSU-155 at a final concentration of 5 µM for 3 days when bacterial load was estimated by fluorescent pixel count. \*\*\**P* = 0.0002 Mann-Whitney test, unpaired, two tailed.

## DISCUSSION

The urgent demand for bactericidal drugs targeting Mabs infections persists, driven by the limited efficacy of current treatments and the ability of the pathogen to evade antibiotic action through intrinsic resistance mechanisms and its lipid-rich cell envelope that prevents drug penetration. Prolonged therapeutic regimens to treat Mabs infections/diseases, often spanning months to years, result in low success rates^5,6^. Repurposing anti-tuberculosis drugs is a promising strategy due to homologous targets shared between mycobacteria, which allows for potential cross-species efficacy^46^. However, despite significant advances in drug discovery, clinical approval for recently discovered anti-TB drugs remains elusive, highlighting the challenges of translating preclinical successes into effective treatments for NTM diseases. A recent review article highlights the importance of dedicated research and development programs aimed specifically at discovering novel therapeutics to address the unique challenges posed by Mabs-PD^46^. Cell wall inhibitors are typically bactericidal, as they disrupt the synthesis or integrity of the bacterial cell envelope, leading to cell death. Clinically, these inhibitors have been pivotal in treating a wide range of infections, with β-lactam antibiotics being a prime example. Their effectiveness underscores the importance of identifying potent cell wall inhibitors for Mabs, a pathogen with a robust and impermeable cell envelope that contributes to its intrinsic resistance^4^. Targeting this critical vulnerability may offer a valuable strategy for combating these infections. Toward this goal, we used a simple ATP-based assay to provide a practical approach for identifying cell wall inhibitors in Mabs. Building on previous studies showing that cell wall inhibitors cause an apparent increase in ATP in the BacTiter-Glo^TM^ assay^17,47,48^, we found that this surge is an artifact of improved lysis, and developed a whole-cell assay exploiting this for rapid assessment of compound effects on the mycobacterial cell wall (Mulholland et al., bioRxiv 2025.10.24.684413). Here, we optimized this approach for Mabs, revealing several β-lactam inhibitors and a novel naphthalene hit that we further optimized and characterized as an MmpL3 inhibitor. MmpL3, a transporter essential for mycolic acid transport and cell envelope integrity, has been validated as a promising drug target, initially in *M. tuberculosis* and subsequently in Mabs^20,30,36,49–54^. Potent *M. tuberculosis* MmpL3 inhibitors, such as SQ109, AU1235, and BM212, have demonstrated variable or limited activity against Mabs, underscoring the need for compounds specifically optimized for this pathogen^20,30,51^. The naphthalene scaffold described in this study and exemplified by GM47-1, addresses this gap as the first MmpL3 inhibitor specifically optimized for Mabs. The lead molecule demonstrated robust activity against clinical isolates, including drug-resistant strains, and maintained efficacy under intracellular conditions where existing antibiotics often fail. Importantly, its analog MSU-155 also exhibited significant potency in the zebrafish embryo infection model, a well-established system that recapitulates key aspects of Mabs pathogenesis^11,23,42,43,45^. As Mabs-PD treatment currently lacks a defined drug regimen, clinical management typically involves prolonged multidrug therapy combining macrolides with injectable β-lactams such as imipenem or cefoxitin. Previous studies have shown that MmpL3 inhibitors, particularly indole carboxamides, synergize with peptidoglycan-targeting β-lactams against Mabs^20^ as well as *M. tuberculosis*^41^. This synergy likely results from complementary mechanisms of action that affect distinct layers of the mycobacterial cell envelope. In our study, GM47-1 also exhibited synergistic bactericidal activity in combination with β-lactams, further supporting the potential of MmpL3 inhibitors for use in designing more effective combination therapies for Mabs-PD. The primary bactericidal effect of GM47-1 arises from its inhibition of MmpL3, disrupting the transport of mycolic acids necessary for cell envelope synthesis. This is evidenced by the accumulation of TMM and the concomitant depletion of TDM, consistent with the established consequences of MmpL3 inhibition^20,34^. This disruption compromises cell envelope integrity, a critical vulnerability for the pathogen. Genetic validation further supports this mechanism, as escape mutants with specific substitutions in the MmpL3 protein exhibit resistance to GM47-1, reinforcing MmpL3 as the primary target. While some mutations overlap with those observed for other inhibitors^20,30^, the presence of distinct substitutions suggests unique drug-target interactions specific to this compound. In addition, partial silencing of *mmpL3* expression in an *mmpL3* conditional knockdown mutant was associated with increased susceptibility to GM47-1, further validating GM47-1 as a potent MmpL3 inhibitor. In addition to its effect on cell wall synthesis, GM47-1 induces respiratory uncoupling, as evidenced by accelerated oxygen consumption. While previous studies have reported uncoupling effects of MmpL3 inhibitors in *M. tuberculosis*^55^, our work provides a more detailed characterization by linking the reduction in intracellular ATP to increased oxygen consumption, primarily mediated by the cyt-*bcc:aa_3_* branch. These findings underscore the impact of GM47-1 on bacterial respiration and energy homeostasis. Additionally, GM47-1 induces ATP leakage into the extracellular milieu, a previously unreported effect among MmpL3 inhibitors, supporting its ability to compromise cell envelope integrity. The apparent ATP increase observed with GM47-1 in the BacTiter-Glo^TM^ assay was shown to be an assay artifact resulting from this leakage, rather than a true physiological rise in intracellular ATP. Consistent with our accompanying manuscript (see Mulholland et al., bioRxiv 2025.10.24.684413), these findings suggest that similar ATP elevations reported for other cell wall inhibitors may also reflect assay-specific artifacts^56,57^ and underscore the importance of validating ATP-based measurements with complementary methods. Together, our findings establish a practical and mechanistically informative assay platform for identifying cell-wall targeting agents in Mabs, addressing a critical bottleneck in early-stage drug discovery. By applying this approach, we uncovered a naphthalene-based MmpL3 inhibitor series exemplified by GM47-1, which not only disrupts cell envelope biogenesis but also impairs bacterial energy metabolism through respiratory uncoupling and ATP leakage. These dual activities underscore the therapeutic promise of GM47-1 and its derivatives as next-generation agents for the treatment of Mabs-PD, a disease with limited current options and high unmet clinical need.

## MATERIALS AND METHODS

### Bacterial strains, growth, and viability

All mycobacteria strains were cultured in Middlebrook 7H9 medium (Becton Dickson and Company Limited, USA) supplemented with 0.05% Tween 80, 0.5% glycerol, and ADS enrichment (bovine serum albumin, D-glucose, and NaCl), or on Middlebrook 7H10 agar (Becton Dickson and Company Limited, USA) supplemented with OADC. The Mabs CIP104536^T^ strain (ATCC 19977) and all clinical isolates used in this study were kindly gifted by Dr Sung Jae Shin and Dr Won-Jung Koh of Samsung Medical Centre, Seoul, South Korea. The luminescent Mabs strain mDB158 expressing the lux13 operon^37^ derived from Mabs CIP104536^T^ was a kind gift from Prof Daniel Barkan from Koret School of Veterinary Medicine, The Hebrew University of Jerusalem. The MabsΔ*cydABDC* and MabsΔ*qcrCAB* strains used in this study were constructed in a previous study^39^. Recombinant *M. abscessus* CIP104536^T^ strain expressing tdTomato was grown in the presence of 500 µg/ml hygromycin^42,43^. Prior to the start of all experiments, replicating cultures were harvested in the logarithmic phase, washed to remove glycerol from the media, and diluted to the specified cell density according to different experiments. The selection marker drug Zeocin (20 μg/mL) was used for maintaining knockout strains as required. BDQ was purchased from Cellagen Technology LLC, USA.

### ATP assay validation and screening of LOPAC^®1280^ library

ATP quantification of whole-cell cultures was performed using the BacTiter-Glo^TM^ Microbial Cell Viability Assay (Promega, USA), optimized for dual-purpose cell wall/bioenergetics screening in Mabs. For assay validation, cefoxitin and BDQ were used as control drugs for increased ATP release and cellular ATP depletion, respectively. DMSO at a final concentration of 1% served as baseline control. The LOPAC^®1280^ library (Sigma-Aldrich, USA), consisting of 1,280 pharmacologically active compounds, was screened against Mabs Δ*cydABDC* at a single concentration of 40 μM. The drugs were dissolved to a concentration of 4 mM in U-bottom 96-well plates. Then, 1 μL of each drug solution from these intermediate plates was transferred into a white 96-well plate to achieve the 40 µM final drug concentration. Exponentially grown culture was washed twice and resuspended at an OD_600_ of 0.05. A total of 100 μL of the prepared culture was inoculated into each assay well. The plates were then placed in an air-tight box and incubated at 32 °C in an incubator for 4 h. For dose response assay, drug compounds were added at 100× concentration in 1 µL volume into each well of a white 96-well plate. The DMSO concentration was maintained at 1% across all wells. Subsequently, a total volume of 100 µL culture was aliquoted into each well, and the plates were incubated at 32 °C for 4 h. After incubation, 100 μL (1:1) of BacTiter-Glo^TM^ reagent was added to each well to measure intracellular ATP for 10 min using BioTek Cytation 3 Cell Imaging Multiple-mode reader. Luminescence readings were analyzed using GraphPad PRISM to compare the ATP content of each sample to the baseline control. BDQ and 1% DMSO was used as a positive and negative (baseline) control in each screening plate.

### MIC determination

Mabs strains were grown to mid-log phase in Middlebrook 7H9 broth medium supplemented with 0.2% glycerol, 0.05% Tween 80, and 10% ADS supplement and washed twice in Middlebrook 7H9 broth medium supplemented with 0.05% Tween 80 and 10% ADS supplement. BDQ or compounds were dissolved in 90% DMSO, and cefoxitin in water. Drugs were 2-fold serially diluted and spotted in 96-well F-bottom plates at 100× final concentration. The culture inoculum was prepared from washed cells resuspended at a density of OD_600_ 0.005, and a total of 200 μL was added to each well. Plates were incubated at 32 °C for 2 days, except MabsΔ*qcrCAB*, which was incubated for 5 days due to its slow growth rate. Growth inhibition (MIC_50_) was determined by measuring absorbance at 600 nm (OD_600_) using a BioTek Cytation-3 Cell Imaging Multiple-mode reader. MIC_50_ was calculated using nonlinear curve fitting in GraphPad. For agar MIC determination, GM47-1 was supplemented in 7H10 agar. In quadrant petri plate a total of 5 mL of 7H10 agar without GM47-1 and with 1, 2, 4 and 8 µM of GM47-1 was poured in individual quadrant and allowed to solidify. A total of approximately 500-1,000 CFU of Mabs were spread on each plate and incubated for 4 days. Agar MIC was defined as the concentration which prevented the appearance of visible colonies.

### Radiolabeling and TLC analysis

Mabs cells were radiolabeled with [1-^14^C]-acetate (final concentration 0.2 µCi/mL; Perkin Elmer NEC084A001MC) for 2 h in fresh 7H9-glycerol medium with 0.05% tyloxapol and then harvested by centrifugation at 4,000 x *g* for 3 min. For analyses of extractable lipids and mycolates released from mAGP, the labeled cells were lysed in 750 μL chloroform:methanol (2:1, v/v) in a 2 mL tube by sonication in a water bath. The sonication was performed three times, each 30 s with 5 s vortexing in between. 250 µL methanol and 350 μL ultrapure water were then added into the tube, which was subjected to centrifugation at 10,000 x *g* for 2 min to allow phase separation. The bottom organic phase containing extractable lipids was collected and dried in a fume hood overnight. The top layer with insoluble materials was dried using a SpeedVac concentrator (Eppendorf). The pellet was resuspended in 400 μL of 13% (w/v) tetrahydrobutylammonium (Millipore, Sigma) and heated at 95 °C for 2 h to release MAs from the mAGP complex. After the samples cooled down to room temperature, the mixture was acidified with concentrated HCl. Free MAs were extracted twice with 500 µL hexane each, pooled and dried in a fume hood overnight. Both the extractable lipids and released MAs were dissolved in 50 µL chloroform:methanol (2:1, v/v) for analysis by TLC. For total FAME analysis, the labeled cells were resuspended in 400 µL of 13% (w/v) tetrahydrobutylammonium (Millipore, Sigma) before the whole mixture was heated at 95 °C for 2 h. After the samples cooled down to room temperature, 200 µL ultrapure water, 400 µL dichloromethane, and 30 µL iodomethane (Millipore, Sigma) were sequentially added. The mixture was left on an orbital shaker for 1 h to allow ester formation. The resulting mixture was separated into two phases by centrifugation at 5,000 x *g* for 2 min. The bottom organic layer was collected and dried in a fume hood overnight. The FAMEs were dissolved in 50 µL chloroform for subsequent TLC analyses. Lipids were spotted on silica gel 60 F_254_ glass TLC plates (Millipore, Sigma), which were developed once with a chloroform:methanol:water (30:8:1, v/v/v) solvent system for total lipids and free MAs, or thrice with hexane:ethyl acetate (95:5, v/v) for FAMEs in a glass chamber. After development, the TLC plate was completely dried in a fume hood for 1 h. It was then exposed to a storage phosphor screen for 2 days followed by visualization by phosphor imaging (Storm, GE Healthcare).

### Mutation frequency and isolation of resistant mutants

Mabs culture was harvested at the exponential phase and resuspended to a final cell density of approximately 3 × 10^9^ CFU/mL. 7H10 agar was supplemented with 2-fold serial dilutions of GM47-1 to achieve final concentrations of 4-40 µM. Each agar plate was inoculated with 200 µL of culture and subsequently incubated at 37°C for at least 6 days. Individual colonies appearing on agar plates were isolated and verified for their resistance profile by performing *in vitro* MIC and ATP assays.

### Whole genome sequencing and analysis

Genomic DNA was isolated and purified using the Quick-DNA Fungal/Bacterial Miniprep Kit (Zymo Research Corp.). For MiSeq, a library was prepared from 1 µg of genomic DNA using the Illumina PCR-fee kit with TruSeq DNA CD indexes (dual barcoded). Sequencing was performed using MiSeq with 2×300 bp sequencing. Library preparation and whole-genome sequencing were performed at the Singapore Centre for Environmental Life Sciences Engineering at Nanyang Technological University, Singapore. Raw reads obtained were checked using FastQC (version 0.11.9) and adapters trimmed using BBDuk from BBMap tools (version 39.79)^58^. The trimmed reads were then used as input into Snippy (version 4.6.0) for SNP identification against the *Mycobacterium abscessus* ATCC 19977 (NC_010397.1) reference genome^59^.

### Bacterial viability assay

An exponentially growing culture of Mabs was washed and resuspended in fresh 7H9-ADS media. The mycobacterial inoculum was adjusted to an OD_600_ of 0.005 in 7H9-ADS medium. One mL of bacterial culture was added to the wells of a 24-well plate containing pre-aliquoted test compound(s). Plates were incubated at 32 °C. The bacterial culture from each well was then serial diluted and plated on 7H10-OADC agar plates that were incubated at 32 °C for 4-6 days to determine CFUs.

### Methylene blue assay

A qualitative assay to demonstrate changes in oxygen consumption was adapted from previously described method^15,24^. Culture was washed twice and resuspended at a density of OD_600_ 0.3 followed by addition of methylene blue at a final concentration of 0.001% (v/v). Two-milliliter screw-cap glass vials were filled with 1.8 mL of Mabs in the presence of the test compound and BDQ was used alongside as positive control drug. Tubes were tightly sealed and incubated at 32 °C inside an anaerobic jar with an anaerobic sachet (AnaeroPack-Anaero, MCG). For the hypoxic experiments, glycerol was omitted.

### Oxygen consumption rate measurements

Oxygen consumption rates (OCR) were measured using the Seahorse XFe96 analyzer. Agilent Seahorse cell culture microtiter plates were coated with 22.4 µg/mL Cell Tak (Corning®). An exponentially growing culture of Mabs was washed and adjusted to a concentration of OD_600_ 0.2 in 90% diluted 7H9+ADS. Diluted 7H9+ADS was prepared by diluting it with addition of 90% volume of PBST (Phosphate Buffered Saline containing 0.05% Tween 80). 50 µL of culture was added to each well and centrifuged to adhere the bacteria to the cell culture microplate (Agilent). Oxygen consumption rates (OCR) of Mabs were measured using the Seahorse XFe96 Analyzer (Agilent). The oxygen consumption rate (OCR) was derived from 4 min of continuous measurements. Each data point was interspaced with 2 min of continuous mixing. Compounds tested were injected in real-time through the injection port. All data were analyzed using the Wave Desktop 10.1.2, as well as the GraphPad Prism 10 software.

### Intracellular killing

THP-1 (ATCC^®^ TIB-202™) cells were seeded in RPMI (supplemented with FBS, HEPES, pyruvate and NEAA) and incubated at 37 °C (5 % CO_2_). Once the confluency in the flask reached 80-90%, cells were harvested by centrifugation at 1500 rpm for 5 min and re-suspended in fresh media. For differentiation of monocytes to macrophages, THP-1 monocyte cells were treated with 200 nM phorbol myristate acetate (PMA) and were distributed at a density of 2 × 10^5^ cells/well in 24-well plates. After 24 h of differentiation, PMA was washed with fresh media, followed by 48-52 h recovery for complete differentiation to macrophages. Surface adhered macrophage monolayers were washed with pre-warmed RPMI media and infected with Mabs at a multiplicity of infection of 1:2 for 60 min. Pre-warmed complete RPMI medium with or without the test drugs was added. GM-47 was used at 2-6 ×MIC_50_, alongside BDQ (0.5 µM), and clarithromycin (10 µM) were used as positive drug controls. Macrophages were lysed using 0.05% SDS to determine mycobacterial viability post-treatment by plating for CFU determination on 7H10-OADC agar plates.

### *In vivo* efficacy using zebrafish larvae model of infection

Zebrafish embryo production was approved by the A*STAR IACUC under protocol 251900. Adult zebrafish (*Danio rerio*) were spawned to fertilize eggs by natural spawning. Fertilized eggs were raised in E3 media supplemented with methylene blue (0.00005%) at 28°C for one day. Embryos were dechorionated with 1 mg/mL of pronase and subsequently raised in 45 µg/mL 1-phenyl 2-thiourea (PTU, Sigma-Aldrich) from one day post fertilization (dpf). Two days post-fertilization embryos were anesthetized with 160 µg/mL tricaine and microinjected with 1500 CFU Mabs (rough morphotype) expressing tdTomato from the pTEC27 plasmid (pMSP12::tdTomato, Addgene plasmid # 30182, gift from Lalita Ramakrishnan) into the caudal vein to generate an acute infection. Infected embryos were recovered into E3 media supplemented with PTU and drug/vehicle as appropriate in 10 mL volume in 6-well plates and incubated at 32 °C for 3 days. Anesthetized larvae were imaged by epifluorescence microscopy under consistent exposure and camera settings within each experiment. Bacterial fluorescent pixel count was carried out in ImageJ as previously described^44^.

### Estimation of extracellular ATP

To estimate ATP levels in the extracellular milieu, cultures were treated as described for the ATP assay, with the exception that the assay was performed in 12-well plates with a total volume of 2 mL/well. At the end of 4 h treatment with drugs, samples were centrifuged, and the supernatant was collected. Cell pellets were resuspended to the same volume. The supernatant was additionally passed through a syringe filter of 0.2 µM pore size (Acrodisc®, Pall Corporation) to remove any cells or debris. A total volume of 100 µL per well was transferred to a white 96-well plate. Clear supernatant and resuspended cells were used for ATP quantification in triplicate. The experiment was performed with three technical replicates and independently repeated at least once.

### Cytotoxicity determination

HepG2 (ATCC® HB-8065™) was maintained on Dulbecco’s Modified Eagle Medium (DMEM) (Cat No. 11965-092) supplemented with 10% Fetal Bovine Serum (FBS) and 1% penicillin-streptomycin (Sigma). The cells were cultured in DMEM supplemented with glucose for at least one week prior to cytotoxicity assay. The cells monolayer was washed with PBS, then treated with 0.05% Tryp-sin-EDTA for 5 min. Fresh DMEM medium was then added at 4X the volume to inactivate trypsin activity. The cells were then centrifuged (300 × g for 5 min) and resuspended in fresh media before being adjusted to a density of 10,000 cells/well in a 96-well F-bottom plate. The cells were then incubated in the plates for 24 h for adhesion and incubated with test compounds for 48 hours before 10 μL of CellTiter 96® AQueous One Solution (Promega) was added to each well. The plates were placed back in the CO_2_ incubator at 37 °C for 2-6 h and read when positive control wells yielded an absorbance reading of 0.8-1.0 at 490 nm. All experiments were conducted in technical triplicates and repeated once.

### General chemistry

All anhydrous solvents, reagent grade solvents for chromatography and starting materials were purchased from either Aldrich Chemical Co. (Milwaukee, WI) or Fisher Scientific (Suwanee, GA) unless otherwise noted. General methods of compound purification involved the use of silica (SiO_2_) purchased from Sorbtec (Norcross, GA) and/or recrystallization. The reactions were monitored by TLC on precoated Merck 60 F_254_ silica gel plates and visualized using UV light (254 nm). All compounds were analyzed for purity by HPLC and characterized by ^1^H and ^13^C NMR spectroscopy using a Bruker DPX Avance I NMR Spectrometer (300MHz) or Bruker Ascend AvanceNeo Spectrometer (actively shielded) (400MHz). Chemical shifts are reported in ppm (δ) relative to the residual solvent peak in the corresponding spectra; chloroform δ 7.27 and δ 77.23, methanol δ 3.31 and δ 49.00 and coupling constants (J) are reported in hertz (Hz) (where, s = singlet, bs = broad singlet, d = doublet, dd = double doublet, bd = broad doublet, ddd = double doublet of doublet, t = triplet, tt – triple triplet, q = quartet, m = multiplet) and analyzed using MestReC NMR data processing. 19F NMR spectra were run without a standard and are uncorrected. Mass spectra values are reported as *m/z*.

### Chemical synthesis

Detailed synthetic methodology of all compounds is summarized in Supplementary Materials (M1-16).

## Supporting information

Supplementary information (Synthesis, Figures and Tables)

## ACKNOWLEDGMENTS

This work was supported by the National Research Foundation (NRF) Singapore under its Investigatorship Program (grant NRF-NRFI06-2020-0004) to K.P. M.B and C.V.M were supported by NIH grants R01 AI173328 and R01 AI175972. G.M. was supported by grant R37AI054193 by the National Institute of Health (NIH), United States. Funding for the Montana State Mass Spectrometry Facility (RRID: SCR_012482) used in this publication was made possible in part by the MJ Murdock Charitable Trust, the National Institute of General Medical Sciences of the National Institutes of Health under Award Numbers P20GM103474 and S10OD28650, and the MSU Office of Research and Economic Development. NMR spectra were recorded at Montana State University on a 400 MHz NMR spectrometer housed in MSU’s NMR Center (RRID:SCR_026334). Financial support for the NMR instruments and operations has been provided in part by the NIH SIG program (1S10RR13878 and 1S10RR026659), the National Science Foundation (NSF-MRI:DBI-1532078; NSF-MRI:CHE-2018388), the Murdock Charitable Trust Foundation (2015066:MNL), and MSU’s Office of Research & Economic Development, and Graduate Education at MSU. L.K. was supported by the Fondation pour la Recherche Médicale (Equipe FRM EQU202103012588). Lipid profiling experiments were funded by Singapore Ministry of Education Academic Research Fund Tier 2 grant MOE-000116 (S.-S.C.). Zebrafish work was supported by Singapore Ministry of Health’s National Medical Research Council under its Individual Research Grant scheme (OFIRG22jul-0081) to S.H.O.

## REFERENCES

1. Johansen, M.D., Herrmann, J.L., and Kremer, L. (2020). Non-tuberculous mycobacteria and the rise of *Mycobacterium abscessus*. Nat. Rev. Microbiol. 18, 392–407. 10.1038/s41579-020-0331-1.

2. Lopeman, R.C., Harrison, J., Desai, M., and Cox, J.A.G. (2019). *Mycobacterium abscessus*: environmental bacterium turned clinical nightmare. Microorganisms 7. 10.3390/microorganisms7030090.

3. Luthra, S., Rominski, A., and Sander, P. (2018). The Role of antibiotic-target-modifying and antibiotic-modifying enzymes in *Mycobacterium abscessus* drug resistance. Front. Microbiol. 9. 10.3389/fmicb.2018.02179.

4. Nessar, R., Cambau, E., Reyrat, J.M., Murray, A., and Gicquel, B. (2012). *Mycobacterium abscessus*: a new antibiotic nightmare. J. Antimicrob. Chemother. 67, 810–818. 10.1093/jac/dkr578.

5. Kwak, N., Dalcolmo, M.P., Daley, C.L., Eather, G., Gayoso, R., Hasegawa, N., Jhun, B.W., Koh, W.J., Namkoong, H., Park, J., et al. (2019). *Mycobacterium abscessus* pulmonary disease: individual patient data meta-analysis. Eur. Respir. J. 54. 10.1183/13993003.01991-2018.

6. Koh, W.-J., Jeong, B.-H., Kim, S.-Y., Jeon, K., Park, K.U., Jhun, B.W., Lee, H., Park, H.Y., Kim, D.H., Huh, H.J., et al. (2016). Mycobacterial characteristics and treatment outcomes in *Mycobacterium abscessus* lung disease. Clin. Infect. Dis. 64, 309–316. 10.1093/cid/ciw724.

7. Maurer, F.P., Bruderer, V.L., Ritter, C., Castelberg, C., Bloemberg, G.V., and Böttger, E.C. (2014). Lack of antimicrobial bactericidal activity in *Mycobacterium abscessus*. Antimicrob. Agents Chemother. 58, 3828–3836. 10.1128/aac.02448-14.

8. Story-Roller, E., Maggioncalda, E.C., and Lamichhane, G. (2019). Select β-lactam combinations exhibit synergy against *Mycobacterium abscessus in vitro*. Antimicrob. Agents Chemother. 63. 10.1128/aac.02613-18.

9. Soroka, D., Dubée, V., Soulier-Escrihuela, O., Cuinet, G., Hugonnet, J.E., Gutmann, L., Mainardi, J.L., and Arthur, M. (2014). Characterization of broad-spectrum *Mycobacterium abscessus* class A β-lactamase. J. Antimicrob. Chemother. 69, 691–696. 10.1093/jac/dkt410.

10. Ferro, B.E., van Ingen, J., Wattenberg, M., van Soolingen, D., and Mouton, J.W. (2015). Time-kill kinetics of antibiotics active against rapidly growing mycobacteria. J. Antimicrob. Chemother. 70, 811–817. 10.1093/jac/dku431.

11. Dubée, V., Bernut, A., Cortes, M., Lesne, T., Dorchene, D., Lefebvre, A.L., Hugonnet, J.E., Gutmann, L., Mainardi, J.L., Herrmann, J.L., et al. (2015). β-lactamase inhibition by avibactam in *Mycobacterium abscessus*. J. Antimicrob. Chemother. 70, 1051–1058. 10.1093/jac/dku510.

12. Lee, B.S., Singh, S., and Pethe, K. (2023). Inhibiting respiration as a novel antibiotic strategy. Curr. Opin. Microbiol. 74, 102327. 10.1016/j.mib.2023.102327.

13. Lindman, M., and Dick, T. (2019). Bedaquiline eliminates bactericidal activity of β-lactams against *Mycobacterium abscessus*. Antimicrob. Agents Chemother. 63. 10.1128/aac.00827-19.

14. Shetty, A., and Dick, T. (2018). Mycobacterial cell wall synthesis inhibitors cause lethal ATP burst. Front. Microbiol. 9. 10.3389/fmicb.2018.01898.

15. Kalia, N.P., Hasenoehrl, E.J., Ab Rahman, N.B., Koh, V.H., Ang, M.L.T., Sajorda, D.R., Hards, K., Gruber, G., Alonso, S., Cook, G.M., et al. (2017). Exploiting the synthetic lethality between terminal respiratory oxidases to kill *Mycobacterium tuberculosis* and clear host infection. Proc. Natl. Acad. Sci. USA 114, 7426–7431. 10.1073/pnas.1706139114.

16. Cook, G.M., Hards, K., Dunn, E., Heikal, A., Nakatani, Y., Greening, C., Crick, D.C., Fontes, F.L., Pethe, K., Hasenoehrl, E., and Berney, M. (2017). Oxidative phosphorylation as a target space for tuberculosis: success, caution, and future directions. Microbiol. Spectr. 5, 5.3.14. doi:10.1128/microbiolspec.TBTB2-0014-2016.

17. Lee, B.S., Hards, K., Engelhart, C.A., Hasenoehrl, E.J., Kalia, N.P., Mackenzie, J.S., Sviriaeva, E., Chong, S.M.S., Manimekalai, M.S.S., Koh, V.H., et al. (2021). Dual inhibition of the terminal oxidases eradicates antibiotic-tolerant *Mycobacterium tuberculosis*. EMBO Mol. Med. 13, e13207. 10.15252/emmm.202013207.

18. Pethe, K., Bifani, P., Jang, J., Kang, S., Park, S., Ahn, S., Jiricek, J., Jung, J., Jeon, H.K., Cechetto, J., et al. (2013). Discovery of Q203, a potent clinical candidate for the treatment of tuberculosis. Nat. Med. 19, 1157–1160. 10.1038/nm.3262.

19. La Rosa, V., Poce, G., Canseco, J.O., Buroni, S., Pasca, M.R., Biava, M., Raju, R.M., Porretta, G.C., Alfonso, S., Battilocchio, C., et al. (2012). MmpL3 is the cellular target of the antitubercular pyrrole derivative BM212. Antimicrob. Agents Chemother. 56, 324–331. 10.1128/aac.05270-11.

20. Dupont, C., Viljoen, A., Dubar, F., Blaise, M., Bernut, A., Pawlik, A., Bouchier, C., Brosch, R., Guérardel, Y., Lelièvre, J., et al. (2016). A new piperidinol derivative targeting mycolic acid transport in *Mycobacterium abscessus*. Mol. Microbiol. 101, 515–529. 10.1111/mmi.13406.

21. Alcaraz, M., Edwards, T.E., and Kremer, L. (2023). New therapeutic strategies for *Mycobacterium abscessus* pulmonary diseases - untapping the mycolic acid pathway. Expert Rev. Anti Infect. Ther. 21, 813–829. 10.1080/14787210.2023.2224563.

22. Lee, B.S., Kalia, N.P., Jin, X.E.F., Hasenoehrl, E.J., Berney, M., and Pethe, K. (2019). Inhibitors of energy metabolism interfere with antibiotic-induced death in mycobacteria. J. Biol. Chem. 294, 1936–1943. 10.1074/jbc.RA118.005732.

23. Dupont, C., Viljoen, A., Thomas, S., Roquet-Banères, F., Herrmann, J.-L., Pethe, K., and Kremer, L. (2017). Bedaquiline inhibits the ATP synthase in *Mycobacterium abscessus* and is effective in infected zebrafish. Antimicrob. Agents Chemother. 61, e01225–01217. doi:10.1128/AAC.01225-17.

24. Kalia, N.P., Singh, S., Hards, K., Cheung, C.Y., Sviriaeva, E., Banaei-Esfahani, A., Aebersold, R., Berney, M., Cook, G.M., and Pethe, K. (2023). *M. tuberculosis* relies on trace oxygen to maintain energy homeostasis and survive in hypoxic environments. Cell Rep. 42, 112444. 10.1016/j.celrep.2023.112444.

25. Chandrasekera, N.S., Berube, B.J., Shetye, G., Chettiar, S., O’Malley, T., Manning, A., Flint, L., Awasthi, D., Ioerger, T.R., Sacchettini, J., et al. (2017). Improved Phenoxyalkylbenzimidazoles with activity against *Mycobacterium tuberculosis* appear to target *QcrB*. ACS Infect. Dis. 3, 898–916. 10.1021/acsinfecdis.7b00112.

26. Moraski, G.C., Seeger, N., Miller, P.A., Oliver, A.G., Boshoff, H.I., Cho, S., Mulugeta, S., Anderson, J.R., Franzblau, S.G., and Miller, M.J. (2016). Arrival of imidazo[2,1-b]thiazole-5-carboxamides: potent anti-tuberculosis agents that target *QcrB*. ACS Infect. Dis. 2, 393–398. 10.1021/acsinfecdis.5b00154.

27. Foo, C.S., Lupien, A., Kienle, M., Vocat, A., Benjak, A., Sommer, R., Lamprecht, D.A., Steyn, A.J.C., Pethe, K., Piton, J., et al. (2018). Arylvinylpiperazine amides, a new class of potent inhibitors targeting *QcrB* of *Mycobacterium tuberculosis*. mBio 9. 10.1128/mBio.01276-18.

28. Roux, A.L., Viljoen, A., Bah, A., Simeone, R., Bernut, A., Laencina, L., Deramaudt, T., Rottman, M., Gaillard, J.L., Majlessi, L., et al. (2016). The distinct fate of smooth and rough *Mycobacterium abscessus* variants inside macrophages. Open Biol. 6. 10.1098/rsob.160185.

29. Zhang, J., Leifer, F., Rose, S., Chun, D.Y., Thaisz, J., Herr, T., Nashed, M., Joseph, J., Perkins, W.R., and DiPetrillo, K. (2018). Amikacin liposome inhalation suspension (ALIS) penetrates non-tuberculous mycobacterial biofilms and enhances amikacin uptake into macrophages. Front. Microbiol. 9, 915. 10.3389/fmicb.2018.00915.

30. Raynaud, C., Daher, W., Johansen, M.D., Roquet-Banères, F., Blaise, M., Onajole, O.K., Kozikowski, A.P., Herrmann, J.-L., Dziadek, J., Gobis, K., and Kremer, L. (2020). Active benzimidazole derivatives targeting the MmpL3 transporter in *Mycobacterium abscessus*. ACS Infect. Dis. 6, 324–337. 10.1021/acsinfecdis.9b00389.

31. Kozikowski, A.P., Onajole, O.K., Stec, J., Dupont, C., Viljoen, A., Richard, M., Chaira, T., Lun, S., Bishai, W., Raj, V.S., et al. (2017). Targeting mycolic acid transport by indole-2-carboxamides for the treatment of *Mycobacterium abscessus* infections. J. Med. Chem. 60, 5876–5888.10.1021/acs.jmedchem.7b00582.

32. Shimao, T. (1987). Drug resistance in tuberculosis control. Tubercle 68, 5–15. 10.1016/0041-3879(87)90014-6.

33. Williams, J.T., and Abramovitch, R.B. (2023). Molecular mechanisms of MmpL3 function and inhibition. Microb. Drug Resist. 29, 190–212.10.1089/mdr.2021.0424.

34. Xu, Z., Meshcheryakov, V.A., Poce, G., and Chng, S.-S. (2017). MmpL3 is the flippase for mycolic acids in mycobacteria. Proc. Natl. Acad. Sci. USA 114, 7993–7998. doi:10.1073/pnas.1700062114.

35. Takayama, K., Wang, C., and Besra, G.S. (2005). Pathway to synthesis and processing of mycolic acids in *Mycobacterium tuberculosis*. Clin. Microbiol. Rev. 18, 81–101. doi:10.1128/cmr.18.1.81-101.2005.

36. Boudehen, Y.M., Tasrini, Y., Aguilera-Correa, J.J., Alcaraz, M., and Kremer, L. (2023). Silencing essential gene expression in *Mycobacterium abscessus* during infection. Microbiol. Spectr. 11, e0283623. 10.1128/spectrum.02836-23.

37. Meir, M., Grosfeld, T., and Barkan, D. (2018). Establishment and validation of *Galleria mellonella* as a novel model organism To study *Mycobacterium abscessus* infection, pathogenesis, and treatment. Antimicrob. Agents Chemother. 62. 10.1128/aac.02539-17.

38. Lamprecht, D.A., Finin, P.M., Rahman, M.A., Cumming, B.M., Russell, S.L., Jonnala, S.R., Adamson, J.H., and Steyn, A.J. (2016). Turning the respiratory flexibility of *Mycobacterium tuberculosis* against itself. Nat. Comm. 7, 12393. 10.1038/ncomms12393.

39. Sorayah, R., Manimekalai, M.S.S., Shin, S.J., Koh, W.-J., Grüber, G., and Pethe, K. (2019). Naturally-occurring polymorphisms in *QcrB* are responsible for resistance to Telacebec in *Mycobacterium abscessus*. ACS Infect. Dis. 5, 2055–2060. 10.1021/acsinfecdis.9b00322.

40. Raynaud, C., Daher, W., Roquet-Banères, F., Johansen, M.D., Stec, J., Onajole, O.K., Ordway, D., Kozikowski, A.P., and Kremer, L. (2020). Synergistic interactions of indole-2-carboxamides and & β-lactam Antibiotics against *Mycobacterium abscessus*. Antimicrob. Agents Chemother. 64, e02548. doi:10.1128/aac.02548-19.

41. Li, W., Sanchez-Hidalgo, A., Jones, V., de Moura, V.C., North, E.J., and Jackson, M. (2017). Synergistic interactions of MmpL3 inhibitors with antitubercular compounds *in vitro*. Antimicrob. Agents Chemother. 61, e02399. 10.1128/aac.02399-16.

42. Bernut, A., Herrmann, J.L., Kissa, K., Dubremetz, J.F., Gaillard, J.L., Lutfalla, G., and Kremer, L. (2014). *Mycobacterium abscessus* cording prevents phagocytosis and promotes abscess formation. Proc. Natl. Acad. Sci. USA 111, E943–952. 10.1073/pnas.1321390111.

43. Bernut, A., Nguyen-Chi, M., Halloum, I., Herrmann, J.L., Lutfalla, G., and Kremer, L. (2016). *Mycobacterium abscessus*-induced granuloma formation is strictly dependent on TNF signaling and neutrophil trafficking. PLoS Pathog. 12, e1005986. 10.1371/journal.ppat.1005986.

44. Matty, M.A., Oehlers, S.H., and Tobin, D.M. (2016). Live imaging of host– pathogen interactions in zebrafish larvae. In Zebrafish: Methods and Protocols, K. Kawakami, E.E. Patton, and M. Orger, eds. (Springer New York), pp. 207–223. 10.1007/978-1-4939-3771-4_14.

45. Bernut, A., Moigne, V.L., Lesne, T., Lutfalla, G., Herrmann, J.-L., and Kremer, L. (2014). *In vivo* assessment of drug efficacy against *Mycobacterium abscessus* using the embryonic zebrafish test system. Antimicrob. Agents Chemother. 58, 4054–4063. doi:10.1128/aac.00142-14.

46. Ganapathy, U.S., and Dick, T. (2022). Why matter matters: fast-tracking *Mycobacterium abscessus* drug discovery. Molecules 27. 10.3390/molecules27206948.

47. Rao, S.P.S., Alonso, S., Rand, L., Dick, T., and Pethe, K. (2008). The protonmotive force is required for maintaining ATP homeostasis and viability of hypoxic, nonreplicating *Mycobacterium tuberculosis*. Proc. Natl. Acad. Sci. USA 105, 11945–11950. 10.1073/pnas.0711697105.

48. Hunter, D.M., and Lim, D.V. (2010). Rapid detection and identification of bacterial pathogens by using an ATP bioluminescence immunoassay. J. Food Prot. 73, 739–746. 10.4315/0362-028x-73.4.739.

49. Lun, S., Guo, H., Onajole, O.K., Pieroni, M., Gunosewoyo, H., Chen, G., Tipparaju, S.K., Ammerman, N.C., Kozikowski, A.P., and Bishai, W.R. (2013). Indoleamides are active against drug-resistant *Mycobacterium tuberculosis*. Nat. Comm. 4, 2907. 10.1038/ncomms3907.

50. Li, W., Obregón-Henao, A., Wallach, J.B., North, E.J., Lee, R.E., Gonzalez-Juarrero, M., Schnappinger, D., and Jackson, M. (2016). Therapeutic potential of the *Mycobacterium tuberculosis* mycolic acid transporter, MmpL3. Antimicrob. Agents Chemother. 60, 5198–5207. 10.1128/aac.00826-16.

51. Tahlan, K., Wilson, R., Kastrinsky, D.B., Arora, K., Nair, V., Fischer, E., Barnes, S.W., Walker, J.R., Alland, D., Barry, C.E., 3rd, and Boshoff, H.I. (2012). SQ109 targets MmpL3, a membrane transporter of trehalose monomycolate involved in mycolic acid donation to the cell wall core of *Mycobacterium tuberculosis*. Antimicrob. Agents Chemother. 56, 1797–1809. 10.1128/aac.05708-11.

52. Degiacomi, G., Benjak, A., Madacki, J., Boldrin, F., Provvedi, R., Palù, G., Kordulakova, J., Cole, S.T., and Manganelli, R. (2017). Essentiality of mmpL and impact of its silencing on *Mycobacterium tuberculosis* gene expression. Sci. Rep. 7, 43495. 10.1038/srep43495.

53. McNeil, M.B., and Cook, G.M. (2019). Utilization of CRISPR Interference t validate MmpL3 as a drug target in *Mycobacterium tuberculosis*. Antimicrob. Agents Chemother. 63, e00629. doi:10.1128/aac.00629-19.

54. Franz, N.D., Belardinelli, J.M., Kaminski, M.A., Dunn, L.C., Calado Nogueira de Moura, V., Blaha, M.A., Truong, D.D., Li, W., Jackson, M., and North, E.J. (2017). Design, synthesis and evaluation of indole-2-carboxamides with pan anti-mycobacterial activity. Bioorg. Med. Chem. 25, 3746–3755. 10.1016/j.bmc.2017.05.015.

55. Foss, M.H., Pou, S., Davidson, P.M., Dunaj, J.L., Winter, R.W., Pou, S., Licon, M.H., Doh, J.K., Li, Y., Kelly, J.X., et al. (2016). Diphenylether-modified 1,2-diamines with improved drug properties for development against *Mycobacterium tuberculosis*. ACS Infect. Dis. 2, 500–508. 10.1021/acsinfecdis.6b00052.

56. Montville, T.J., Chung, H.-J., Chikindas, M.L., and Chen, Y. (1999). Nisin A depletes intracellular ATP and acts in bactericidal manner against *Mycobacterium smegmatis*. Lett. Appl. Microbiol. 28, 189–193. 10.1046/j.1365-2672.1999.00511.x.

57. Wang, X., van Beekveld, R.A.M., Xu, Y., Parmar, A., Das, S., Singh, I., and Breukink, E. (2023). Analyzing mechanisms of action of antimicrobial peptides on bacterial membranes requires multiple complimentary assays and different bacterial strains. Biochim. Biophys. Acta 1865, 184160. 10.1016/j.bbamem.2023.184160.

58. Bushnell, B. (2015). BBMap short-read aligner, and other bioinformatics tools.

59. Seemann, T. (2015). Snippy: fast bacterial variant calling from NGS reads.

