## Supplementary information (Synthesis, Figures and Tables) for "A bioluminescence-based chemical screen identifies a bactericidal naphthalene scaffold targeting MmpL3 in *Mycobacterium abscessus*"

### CHEMISTRY METHODS

#### Method M1. Synthesis of GM46-96 (SBI-0087702)

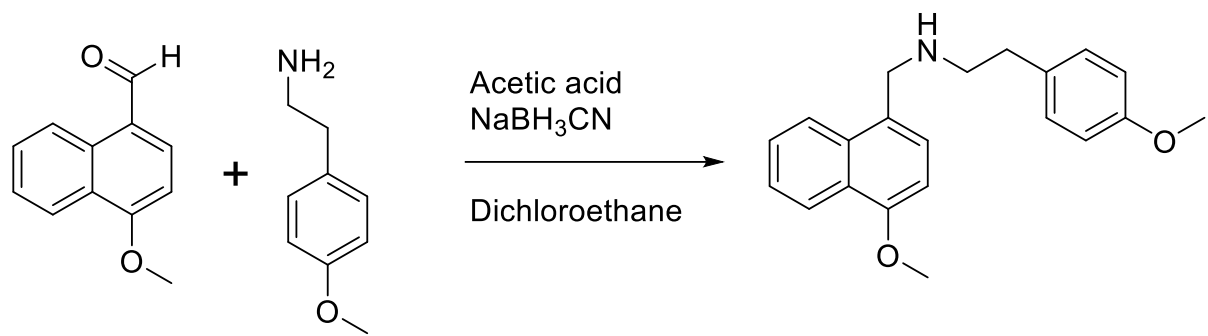

In Rb flask, 4-methoxynaphthalenecarbaldehyde (0.120 g, 0.63 mmol) and 2-(4-methoxyphenyl)ethan-1-amine (0.190 g, 1.26 mmol) were combined in 5 mL anhydrous DCE. Next, acetic acid (0.18 mL, 3.16 mmol) was added (apparent pH~5 when checked by wet pH paper) and the reaction was stirred at RT under argon for 5 h. At which time, NaBH<sub>3</sub>CN (0.061 g, 0.95 mmol) was added, and rapid bubbling was observed. Reaction stirred at room temperature under argon. Once complete by TLC (EtOAc eluent) the reaction was diluted with CH<sub>2</sub>Cl<sub>2</sub> and transferred to a separatory funnel. The organic layer was washed with sat. NaHCO<sub>3</sub> aq. solution (2x), brine, and then organics were dried over Na<sub>2</sub>SO<sub>4</sub>. The drying agent was removed by filtration, and the organics were concentrated down to a clear oil. The residue was purified through a silica gel column, eluting with CH<sub>2</sub>Cl<sub>2</sub> to remove upper running spots (starting materials). Then, the polarity was increased with ethyl acetate from 0% to 25%, N-((4-methoxynaphthalen-1-yl)methyl)-2-(4-methoxyphenyl)ethan-1-amine (**SBI-0087702**, **GM46-96**) was collected. Fractions were concentrated to yield 125 mg (62%) of a white solid. HRMS (ESI-TOF, positive mode) m/z ([M+1]); Anal. Calcd. for C<sub>21</sub>H<sub>24</sub>NO<sub>2</sub>, 322.1802; found 322.1791; RT= 3.31 min. <sup>1</sup>H NMR (300 MHz, CDCl<sub>3</sub>) δ 8.31 – 8.21 (m, 1H), 7.99 – 7.88 (m, 1H), 7.54 – 7.39 (m, 2H), 7.30 (d, J = 7.8 Hz, 1H), 7.15 – 7.06 (m, 2H), 6.85 – 6.75 (m, 2H), 6.71 (d, J = 7.8 Hz, 1H), 4.13 (s, 2H), 3.97 (s, 3H), 3.76 (s, 3H), 2.95 (t, J = 7.1 Hz, 2H), 2.78 (t, J = 7.1 Hz, 2H).

#### Method M2. Synthesis of MSU-06

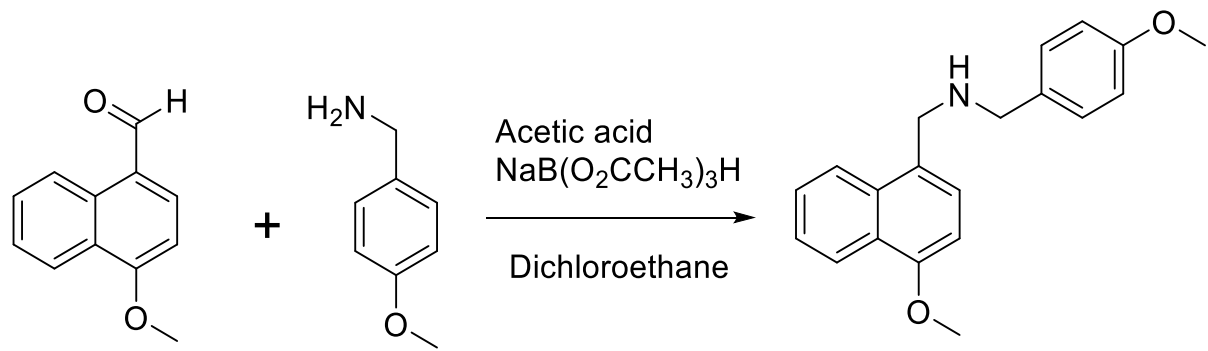

In Rb flask, 4-methoxynaphthalenecarbaldehyde (0.11 g, 0.58 mmol) and 4-methoxyphenylmethylamine (0.81 g, 0.58 mmol) were combined in 5 mL DCE. Next, acetic acid (0.13 mL, 2.32 mmol) was added (apparent pH~5 when checked by wet pH paper) and the reaction was stirred at RT under argon for 1 h. At which time, NaB(O<sub>2</sub>CCH<sub>3</sub>)<sub>3</sub>H (0.184 g, 0.87 mmol) was added, and slow bubbling was observed. Reaction stirred at room temperature under argon. Once complete by TLC (EtOAc eluent) or HPLC, the reaction was diluted with CH<sub>2</sub>Cl<sub>2</sub> and transferred to a separatory funnel. The organic layer was washed with sat. NaHCO<sub>3</sub> aq. Solution (2x), brine, and then organics were dried over Na<sub>2</sub>SO<sub>4</sub>. The drying agent was removed by filtration, and the organics were concentrated down to a clear oil. The residue was purified through a silica gel column, eluting with $\text{CH}_2\text{Cl}_2$ to remove upper running spots (starting materials). Then, the polarity was increased with ethyl acetate from 0% to 50%, and *N*-(4-methoxybenzyl)-1-(4-methoxynaphthalen-1-yl)methanamine (**MSU-06**) was collected. Fractions were concentrated to yield 107 mg (57%) of a yellow oil. Calcd. for $\text{C}_{20}\text{H}_{22}\text{NO}_2$, 308.1651; found 308.1640; RT= 3.28 min. $^1\text{H}$ NMR (300 MHz, MeOD) $\delta$ 8.27 – 8.18 (m, 1H), 7.85 (dd, $J$ = 7.8, 1.7 Hz, 1H), 7.53 – 7.38 (m, 3H), 7.35 (d, $J$ = 7.9 Hz, 1H), 7.31 – 7.21 (m, 2H), 6.94 – 6.76 (m, 3H), 4.04 (s, 2H), 3.97 (s, 3H), 3.77 (s, 3H).

#### Method M3. Synthesis of MSU-21

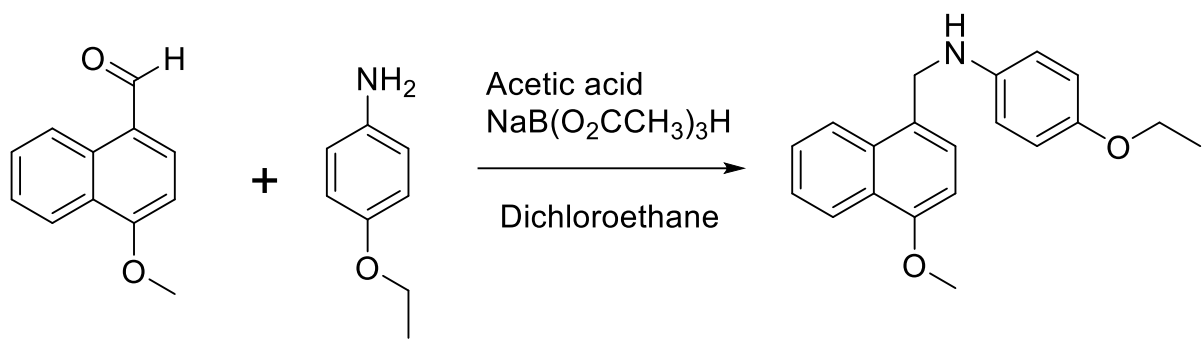

In Rb flask, 4-methoxynaphthalenecarbaldehyde (0.15 g, 0.79 mmol) and 4-ethoxyaniline (0.108 g, 0.79 mmol) were combined in 5 mL DCE. Next, acetic acid (0.18 mL, 3.16 mmol) was added (apparent pH~5 when checked by wet pH paper) and the reaction was stirred at RT under argon for 1 h. At which time, $\text{NaB}(\text{O}_2\text{CCH}_3)_3\text{H}$ (0.251 g, 1.18 mmol) was added, and slow bubbling was observed. Reaction stirred at room temperature under argon. Once complete by TLC (EtOAc eluent) or HPLC, the reaction was diluted with $\text{CH}_2\text{Cl}_2$ and transferred to a separatory funnel. The organic layer was washed with sat. $\text{NaHCO}_3$ aq. Solution (2x), brine, and then organics were dried over $\text{Na}_2\text{SO}_4$. The drying agent was removed by filtration, and the organics were concentrated down to a clear oil. The residue was purified through a silica gel column, eluting with $\text{CH}_2\text{Cl}_2$ to remove upper running spots (starting materials). Then, the polarity was increased with ethyl acetate from 0% to 50%, and 4-ethoxy-*N*-((4-methoxynaphthalen-1-yl)methyl)aniline (**MSU-21**) was collected. Fractions were concentrated to yield 185 mg (72%) of an off-white solid. HRMS (ESI-TOF, positive mode) $m/z$ ($[\text{M}+1]$); Anal. Calcd. for $\text{C}_{20}\text{H}_{22}\text{NO}_2$, 308.1651; found 308.1604; RT= 4.88 min. $^1\text{H}$ NMR (400 MHz, $\text{CDCl}_3$) $\delta$ 8.38 – 8.31 (m, 1H), 8.05 (dd, $J$ = 7.6, 1.7 Hz, 1H), 7.56 – 7.48 (m, 2H), 7.45 (d, $J$ = 7.8 Hz, 1H), 6.88 – 6.81 (m, 2H), 6.78 (d, $J$ = 7.9 Hz, 1H), 6.72 – 6.64 (m, 2H), 4.62 (s, 2H), 4.05 – 3.96 (m, 5H, OCH<sub>3</sub> and OCH<sub>2</sub>), 1.41 (t, $J$ = 7.0 Hz, 3H).

#### Method M4. Synthesis of MSU-19

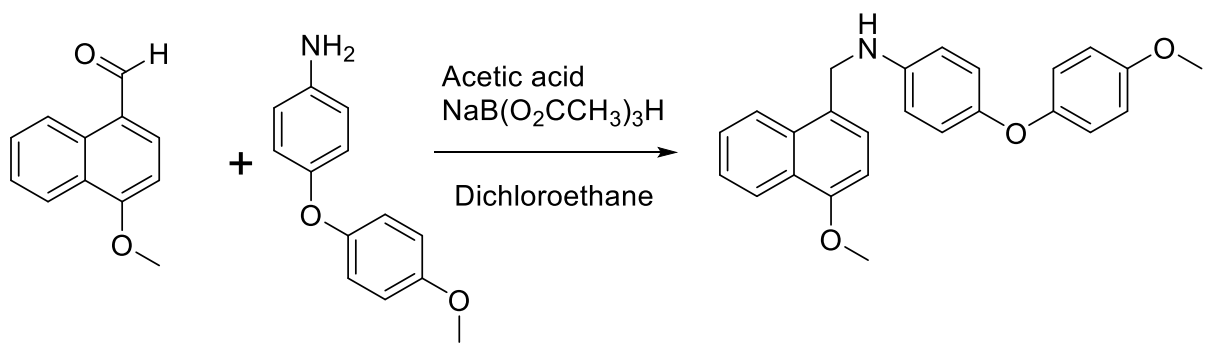

In Rb flask, 4-methoxynaphthalenecarbaldehyde (0.087 g, 0.46 mmol) and 4-(4-methoxyphenoxy)aniline (0.101 g, 0.46 mmol) were combined in 5 mL DCE. Next, acetic acid (0.11 mL, 1.84 mmol) was added (apparent pH~5 when checked by wet pH paper) and the reaction was stirred at RT under argon for 1 h. At which time, $\text{NaB}(\text{O}_2\text{CCH}_3)_3\text{H}$ (0.146 g, 0.69 mmol) was added, and slow bubbling was observed. Reaction stirred at room temperature under argon. Once complete by TLC (EtOAc eluent) or HPLC, the reaction was diluted with $\text{CH}_2\text{Cl}_2$ and transferred to a separatory funnel. The organic layer was washed with sat. $\text{NaHCO}_3$ aq. Solution (2x), brine, and then organics were dried over $\text{Na}_2\text{SO}_4$. The drying agent was removed by filtration, and the organics were concentrated down to a clear oil. The residue was purified through a silica gel column, eluting with $\text{CH}_2\text{Cl}_2$ to remove upper running spots (starting materials). Then, the polarity was increased with ethyl acetate from 0% to 50%, and *N*-((4-methoxynaphthalen-1-yl)methyl)-4-(4-methoxyphenoxy)aniline (**MSU-19**) was collected. Fractions were concentrated to yield 142 mg (76%) of a glassy solid. HRMS (ESI-TOF, positive mode) $m/z$ ($[\text{M}+1]$); Anal. Calcd. for $\text{C}_{25}\text{H}_{23}\text{NO}_3$ 386.1756; found 386.1705. RT= 6.50 min. $^1\text{H}$ NMR (400 MHz, $\text{CDCl}_3$) $\delta$ 8.36 (d, $J$ = 8.2 Hz, 1H), 8.05 (d, $J$ = 8.2 Hz, 1H), 7.55 (p, $J$ = 7.1 Hz, 2H), 7.46 (d, $J$ = 7.8 Hz, 1H), 6.93 (dd, $J$ = 12.3, 8.4 Hz, 4H), 6.86 (d, $J$ = 8.6 Hz, 2H), 6.79 (d, $J$ = 7.8 Hz, 1H), 6.69 (d, $J$ = 8.3 Hz, 2H), 4.64 (s, 2H), 4.04 (s, 3H), 3.81 (s, 3H).

##### Method M5. Synthesis of GM47-1

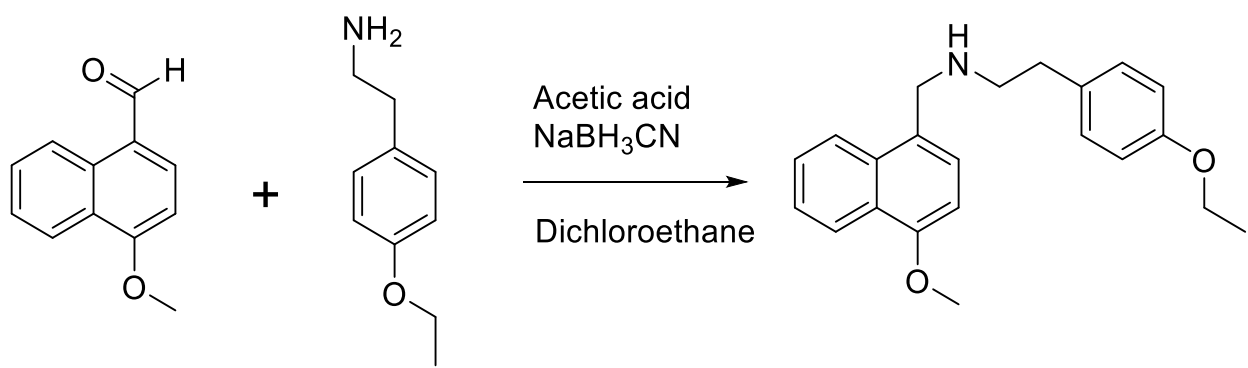

In Rb flask, 4-methoxynaphthalenecarbaldehyde (0.385 g, 2.0 mmol) and 2-(4-ethoxyphenyl)ethanamine (0.410 g, 2.43 mmol) were combined in 15 mL DCE. Next, acetic acid (0.47 mL, 8.1 mmol) was added (apparent pH~5 when checked by wet pH paper) and the reaction was stirred at RT under argon for 1 h. At which time, $\text{NaBH}_3\text{CN}$ (0.127 g, 2.0 mmol) was added, and slow bubbling was observed. Reaction stirred at room temperature under argon. Once complete by TLC (EtOAc eluent) or HPLC, the reaction was diluted with $\text{CH}_2\text{Cl}_2$ and transferred to a separatory funnel. The organic layer was washed with sat. $\text{NaHCO}_3$ aq. Solution (2x), brine, and then organics were dried over $\text{Na}_2\text{SO}_4$. The drying agent was removed by filtration, and the organics were concentrated down to a clear oil. The residue was purified through a silica gel column, eluting with $\text{CH}_2\text{Cl}_2$ to remove upper running spots (starting materials). Then, the polarity was increased with ethyl acetate from 0% to 50%, and 2-(4-ethoxyphenyl)-*N*-((4-methoxynaphthalen-1-yl)methyl)ethan-1-amine (**GM47-1**) was collected. Fractions were concentrated to yield 215 mg (30%) of an off-white solid. HRMS (ESI-TOF, positive mode) $m/z$ ($[\text{M}+1]$); Anal. Calcd. for $\text{C}_{22}\text{H}_{26}\text{NO}_2$, 336.1964; found 336.1948 ; RT= 4.85 min. $^1\text{H}$ NMR (400 MHz, MeOD) $\delta$ 8.30 – 8.22 (m, 1H), 7.86 (dd, $J$ = 7.3, 2.0 Hz, 1H), 7.48 (dq, $J$ = 8.4, 6.8, 1.6 Hz, 2H), 7.37 (d, $J$ = 7.8 Hz, 1H), 7.13 – 7.05 (m, 2H), 6.86 (d, $J$ = 7.8 Hz, 1H), 6.84 – 6.77 (m, 2H), 4.13 (s, 2H), 4.01 (s, $J$ = 6.6 Hz, 3H), 4.00 (q, $J$ = 7.0 Hz, 2H), 2.93 (t, $J$ = 7.3 Hz, 2H), 2.79 (t, $J$ = 7.3 Hz, 2H), 1.38 (t, $J$ = 7.0 Hz, 3H).

##### Method M6. Synthesis of MSU-24

In Rb flask, 4-methoxynaphthalenecarbaldehyde (0.152 g, 0.80 mmol) and (3-ethoxyphenyl)ethanamine (0.135 g, 0.80 mmol) were combined in 5 mL DCE. Next, acetic acid (0.18 mL, 3.2 mmol) was added (apparent pH~5 when checked by wet pH paper) and the reaction was stirred at RT under argon for 1 h. At which time, $\text{NaB}(\text{O}_2\text{CCH}_3)_3\text{H}$ (0.254 g, 1.20 mmol) was added, and slow bubbling was observed. Reaction stirred at room temperature under argon. Once complete by TLC (EtOAc eluent) or HPLC, the reaction was diluted with $\text{CH}_2\text{Cl}_2$ and transferred to a separatory funnel. The organic layer was washed with sat. $\text{NaHCO}_3$ aq. Solution (2x), brine, and then organics were dried over $\text{Na}_2\text{SO}_4$. The drying agent was removed by filtration, and the organics were concentrated down to a clear oil. The residue was purified through a silica gel column, eluting with $\text{CH}_2\text{Cl}_2$ to remove upper running spots (starting materials). Then, the polarity was increased with ethyl acetate from 0% to 50%, and 2-(3-ethoxyphenyl)-*N*-((4-methoxynaphthalen-1-yl)methyl)ethan-1-amine (**MSU-24**) was collected. Fractions were concentrated to yield 158 mg (56%) of a light-yellow oil. HRMS (ESI-TOF, positive mode) $m/z$ ($[\text{M}+1]$); Anal. Calcd. for $\text{C}_{22}\text{H}_{26}\text{NO}_2$, 336.1964; found 336.1966; RT= 3.72 min. $^1\text{H}$ NMR (400 MHz, $\text{CDCl}_3$) $\delta$ 8.26 – 8.16 (m, 1H), 7.92 – 7.81 (m, 1H), 7.47 – 7.32 (m, 2H), 7.28 (d, $J$ = 7.8 Hz, 1H), 7.09 – 6.91 (m, 2H), 6.77 (td, $J$ = 7.4, 1.1 Hz, 1H), 6.69 (s, 1H), 6.65 (d, $J$ = 7.8 Hz, 1H), 4.16 (s, 2H), 3.90 (s, 3H), 3.85 (q, $J$ = 7.0 Hz, 2H), 2.99 – 2.89 (m, 2H), 2.84 (t, $J$ = 7.1 Hz, 2H), 1.22 (t, $J$ = 7.0 Hz, 3H).

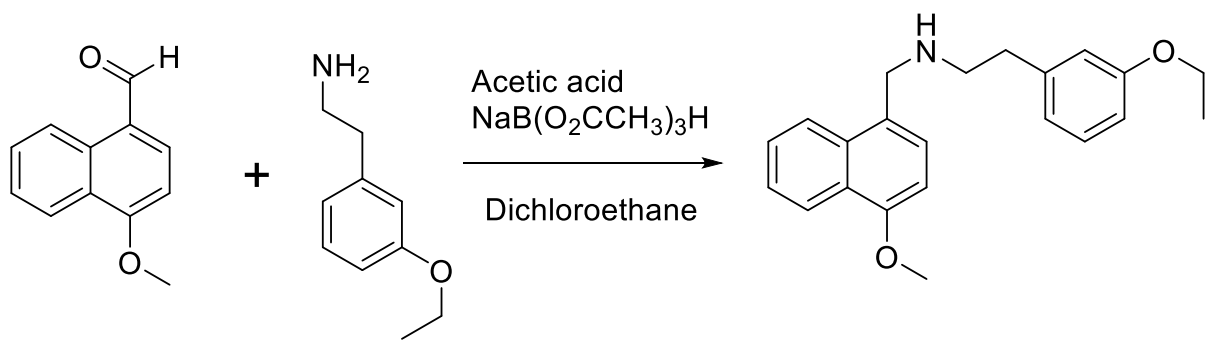

##### Method M7. Synthesis of MSU-26

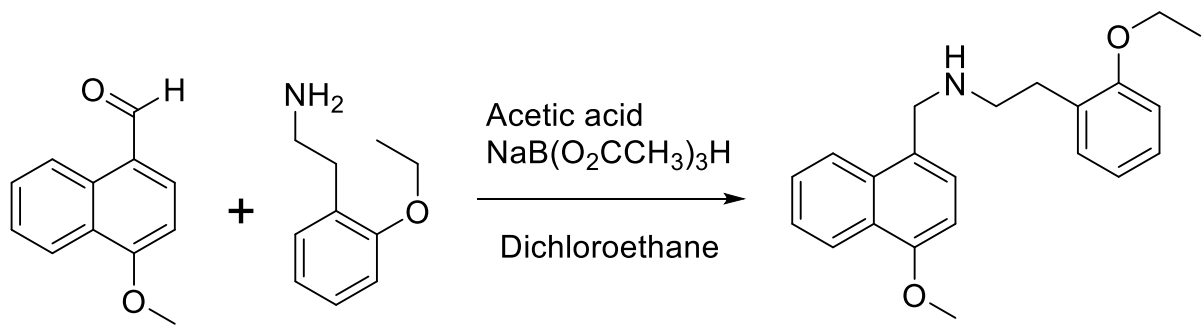

In Rb flask, 4-methoxynaphthalenecarbaldehyde (0.152 g, 0.80 mmol) and (2-(ethoxy)phenyl)ethanamine (0.135 g, 0.80 mmol) were combined in 5 mL DCE. Next, acetic acid (0.18 mL, 3.2 mmol) was added (apparent pH~5 when checked by wet pH paper) and the reaction was stirred at RT under argon for 1 h. At which time, $\text{NaB}(\text{O}_2\text{CCH}_3)_3\text{H}$ (0.254 g, 1.20 mmol) was added, and slow bubbling was observed. Reaction stirred at room temperature under argon. Once complete by TLC (EtOAc eluent) or HPLC, the reaction was diluted with $\text{CH}_2\text{Cl}_2$ and transferred to a separatory funnel. The organic layer was washed with sat. $\text{NaHCO}_3$ aq. Solution (2x), brine, and then organics were dried over $\text{Na}_2\text{SO}_4$. The drying agent was removed by filtration, and the organics were concentrated down to a clear oil. The residue was purified through a silica gel column, eluting with $\text{CH}_2\text{Cl}_2$ to remove upper running spots (starting materials). Then, the polarity was increased with ethyl acetate from 0% to 50%, and 2-(2-ethoxyphenyl)-*N*-((4-methoxynaphthalen-1-yl)methyl)ethan-1-amine (**MSU-26**) was collected. Fractions were concentrated to yield 133 mg (47%) of a yellow oil. HRMS (ESI-TOF, positive mode) $m/z$ ($[\text{M}+1]$); Anal. Calcd. for $\text{C}_{22}\text{H}_{26}\text{NO}_2$, 336.1964; found 336.1966; RT= 3.83 min. $^1\text{H}$ NMR (400 MHz, $\text{CDCl}_3$) $\delta$ 8.20 (dd, $J$ = 7.7, 1.9 Hz, 1H), 7.85 (dd, $J$ = 7.8, 1.8

Hz, 1H), 7.45 – 7.33 (m, 2H), 7.23 (d, $J = 7.8$ Hz, 1H), 7.14 – 7.02 (m, 1H), 6.70 – 6.58 (m, 4H), 4.06 (s, 2H), 3.94 – 3.85 (m, 5H, OCH<sub>3</sub>, OCH<sub>2</sub>), 2.90 (t, $J = 7.2$ Hz, 2H), 2.72 (t, $J = 7.2$ Hz, 2H), 2.65 – 2.55 (m, 2H), 1.30 (t, $J = 7.0$ Hz, 3H).

##### Method M8. Synthesis of MSU-37

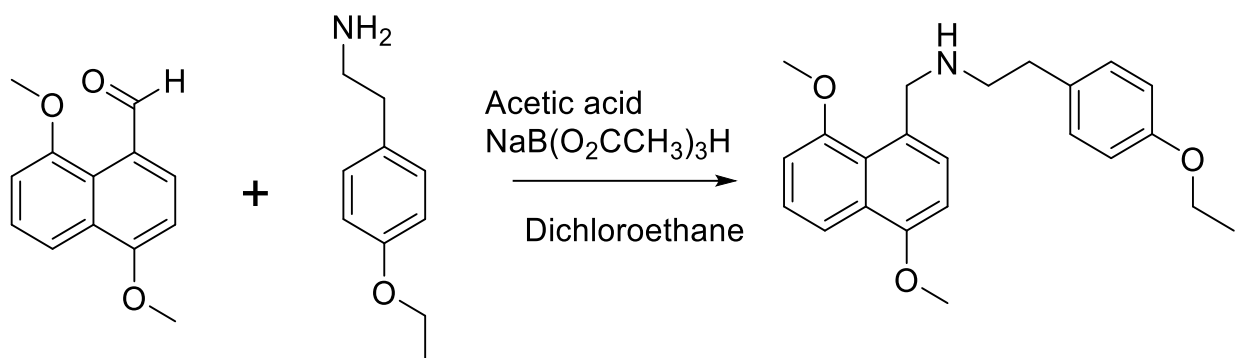

In Rb flask, 4,8-dimethoxy-1-napthaldehyde (0.119 g, 0.54 mmol) and (4-(ethoxy)phenyl)ethanamine (0.91 g, 0.54 mmol) were combined in 5 mL DCE. Next, acetic acid (0.12 mL, 2.2 mmol) was added (apparent pH~5 when checked by wet pH paper) and the reaction was stirred at RT under argon for 1 h. At which time, NaB(O<sub>2</sub>CCH<sub>3</sub>)<sub>3</sub>H (0.171 g, 0.81 mmol) was added, and slow bubbling was observed. Reaction stirred at room temperature under argon. Once complete by TLC (EtOAc eluent) or HPLC, the reaction was diluted with CH<sub>2</sub>Cl<sub>2</sub> and transferred to a separatory funnel. The organic layer was washed with sat. NaHCO<sub>3</sub> aq. solution (2x), brine, and then organics were dried over Na<sub>2</sub>SO<sub>4</sub>. The drying agent was removed by filtration, and the organics were concentrated down to a clear oil. The residue was purified by recrystallization with hot CH<sub>3</sub>CN and *N*-((4,8-dimethoxynaphthalen-1-yl)methyl)-2-(4-ethoxyphenyl)ethan-1-amine (**MSU-37**) (37 mg, 18%) was collected by filtration as an off white solid. HRMS (ESI-TOF, positive mode) $m/z$ ([*M*+1]); Anal. Calcd. for C<sub>23</sub>H<sub>28</sub>NO<sub>3</sub>, 366.2069; found 366.2092; RT= 4.33 min. <sup>1</sup>H NMR (400 MHz, CDCl<sub>3</sub>) $\delta$ 7.85 (dd, $J = 8.4, 1.1$ Hz, 1H), 7.28 (t, $J = 8.1$ Hz, 1H), 7.16 (d, $J = 7.9$ Hz, 1H), 6.96 – 6.88 (m, 2H), 6.75 (d, $J = 7.7$ Hz, 1H), 6.66 (dd, $J = 8.1, 5.6$ Hz, 3H), 4.16 (s, 2H), 3.90 (s, 3H), 3.90 (q, $J = 7.0$ Hz, 2H), 3.67 (s, 3H), 2.75 (t, $J = 6.4$ Hz, 2H), 2.68 (t, $J = 6.4$ Hz, 2H), 1.32 (t, $J = 7.0$ Hz, 3H).

##### Method M9. Synthesis of MSU-43

In a V-vial, 4-methoxynaphthalenecarboxylic acid (0.075 g, 0.37 mmol) was dissolved in 5 ml of anhydrous CH<sub>3</sub>CN and then 2-(4-methoxyphenyl)ethylamine (0.056 g, 0.37 mmol) was added, followed by the EDC-HCl (0.106 g, 0.55 mmol) and DMAP (0.068 g, 0.55 mmol). The reaction vial was sealed and stirred at RT for 12 hours. The reaction mixture was diluted with CH<sub>2</sub>Cl<sub>2</sub>, washed with 10% aqueous NaHCO<sub>3</sub> solution (2x), water, and then 5% acetic acid solution (2x). The organic phase was collected, and then organics were dried over Na<sub>2</sub>SO<sub>4</sub>. The drying agent was removed by filtration, and the organics were concentrated down to a solid. Recrystallized from hot CH<sub>3</sub>CN to give 4-methoxy-*N*-(4-methoxyphenethyl)-1-naphthamide (**MSU-43**) as a white solid (83 mg, 67%). HRMS (ESI-TOF, positive mode) *m/z* ([*M*+1]); Anal. Calcd. for C<sub>21</sub>H<sub>22</sub>NO<sub>3</sub>, 336.1600; found 336.1596; RT= 4.90 min. <sup>1</sup>H NMR (300 MHz, CDCl<sub>3</sub>) δ 8.24 (tt, *J* = 7.1, 2.7 Hz, 2H), 7.56 – 7.40 (m, 3H), 7.22 – 7.12 (m, 2H), 6.90 – 6.79 (m, 2H), 6.70 (d, *J* = 7.9 Hz, 1H), 3.99 (s, 3H), 3.80 – 3.68 (m, 5H, OCH<sub>3</sub> and OCH<sub>2</sub>), 2.91 (t, *J* = 6.8 Hz, 2H).

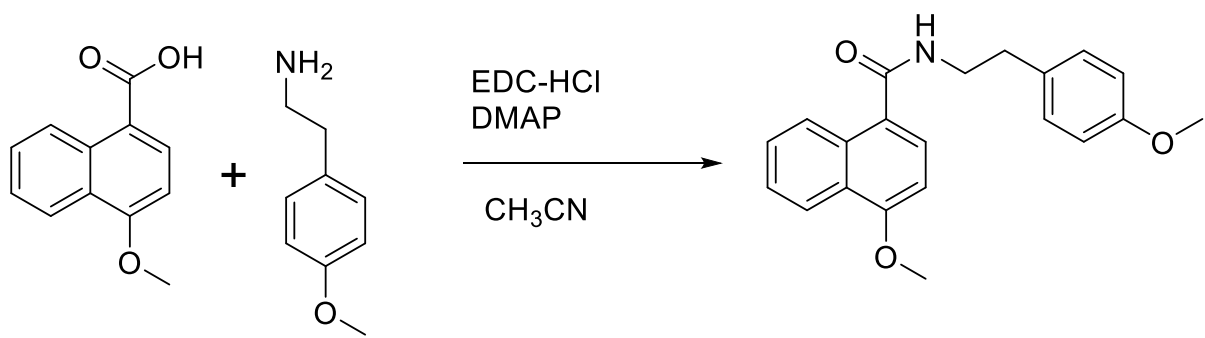

##### Method M10. Synthesis of MSU-134

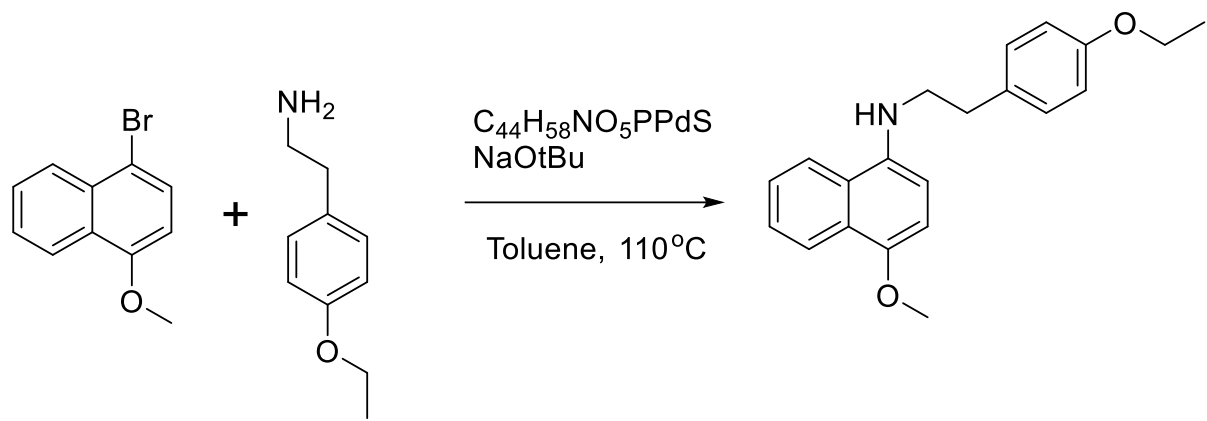

To a sealed tube was combined 1-Bromo-4-methoxynaphthalene (0.133 g, 0.56 mmol), 2-(4-ethoxyphenyl)ethan-1-amine (0.093 g, 0.56 mmol), C<sub>44</sub>H<sub>58</sub>NO<sub>5</sub>PPdS (0.047 g, 0.06 mmol) in 2 mL anhydrous toluene. Argon was bubbled through the solution for 5 minutes, whereupon NaOtBu (0.06 g, 0.68 mmol) was added, sparged for 1 min, sealed, and heated at 110 °C for 48 h. The reaction mixture was filtered over a pad of celite, washed with DCM, and concentrated *in vacuo* to a dark residue. The residue was purified through a silica gel column, eluting with CH<sub>2</sub>Cl<sub>2</sub> to remove upper running spots (starting materials). Then, the polarity was increased with ethyl acetate from 0% to 100%, and *N*-(4-ethoxyphenethyl)-4-methoxynaphthalen-1-amine (**MSU-134**) was collected. Fractions were concentrated to yield 44 mg (23%) of a red rust oil. HRMS (ESI-TOF, positive mode) *m/z* ([*M*+1]); Anal. Calcd. for C<sub>21</sub>H<sub>24</sub>NO<sub>2</sub>, 322.1807; found 322.1805; RT= 5.73 min. <sup>1</sup>H NMR (400 MHz, CDCl<sub>3</sub>) δ 8.31 – 8.25 (m, 1H), 7.89 – 7.80 (m, 2H), 7.58 – 7.46 (m, 2H), 7.19 (d, *J* = 8.6 Hz, 1H), 6.91 – 6.79 (m, 3H), 6.75 (d, *J* = 8.2 Hz, 1H) 4.04 (q, *J* = 7.0 Hz, 2H), 3.98 (s, 3H) 3.52 (t, *J* = 7.2 Hz, 2H), 3.08 (t, *J* = 7.2 Hz, 2H), 1.43 (t, *J* = 7.0 Hz, 3H).

##### Method M11. Synthesis of GM46-72

In Rb flask, 4-methoxynaphthalenecarbaldehyde (0.08 g, 0.42 mmol) and 3-phenylpropylamine (0.114 g, 0.84 mmol) were combined in 5 mL DCE. Next, acetic acid (0.12 mL, 2.1 mmol) was added (apparent pH~5 when checked by wet pH paper) and the reaction was stirred at RT under argon for 1 h. At which time, NaBH<sub>3</sub>CN (0.027 g, 0.42 mmol) was added, and slow bubbling was observed. Reaction stirred at room temperature under argon.

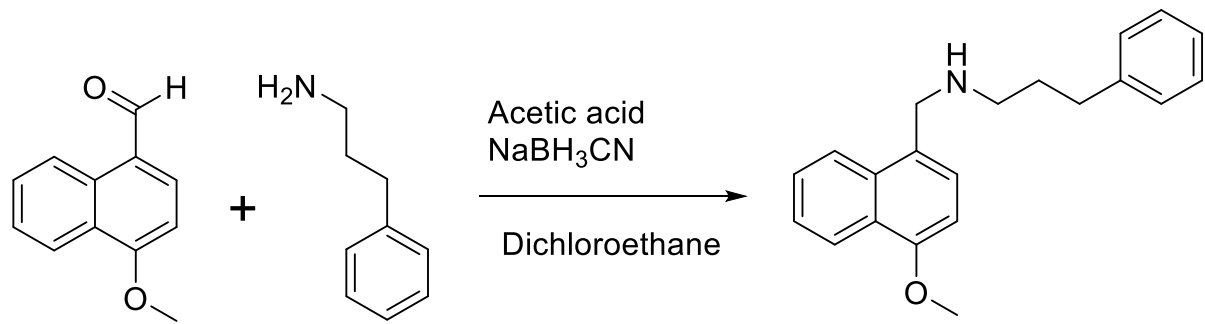

Once complete by TLC (EtOAc eluent) or HPLC, the reaction was diluted with CH<sub>2</sub>Cl<sub>2</sub> and transferred to a separatory funnel. The organic layer was washed with sat. NaHCO<sub>3</sub> aq. solution (2x), brine, and then organics were dried over Na<sub>2</sub>SO<sub>4</sub>. The drying agent was removed by filtration, and the organics were concentrated down to a clear oil. The residue was purified through a silica gel column, eluting with CH<sub>2</sub>Cl<sub>2</sub> to remove upper running spots (starting materials). Then, the polarity was increased with ethyl acetate from 0% to 25%, and *N*-((4-methoxynaphthalen-1-yl)methyl)-3-phenylpropan-1-amine (**GM46-72**) was collected. Fractions were concentrated to yield 42 mg (33%) of an off-white solid. HRMS (ESI-TOF, positive mode) *m/z* ([*M*+1]); Anal. Calcd. for C<sub>21</sub>H<sub>24</sub>NO, 306.1858; found 306.1860; RT= 3.58 min. <sup>1</sup>H NMR (400 MHz, CDCl<sub>3</sub>) δ 8.53 – 8.14 (m, 1H), 7.98 (d, *J* = 8.6 Hz, 1H), 7.60 – 7.37 (m, 3H), 7.27 (d, *J* = 7.9 Hz, 1H), 7.17 – 6.94 (m, 4H), 6.67 (d, *J* = 7.8 Hz, 1H), 4.09 (s, 2H), 3.92 (s, 3H), 2.71 (q, *J* = 7.6 Hz, 2H), 2.59 (t, *J* = 7.7 Hz, 2H), 1.82 (td, *J* = 13.9, 6.5 Hz, 2H).

##### Method M12. Synthesis of GM46-75

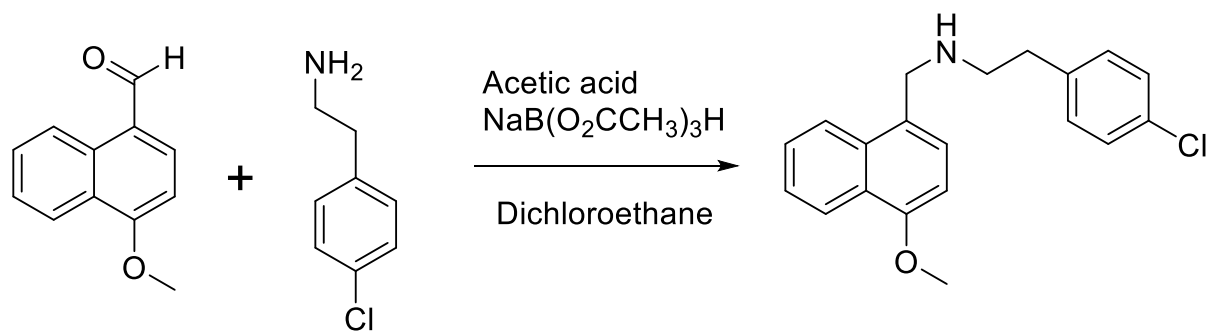

In Rb flask, 4-methoxynaphthalenecarbaldehyde (0.08 g, 0.42 mmol) and 2-(4-chlorophenyl)ethan-1-amine (0.131 g, 0.84 mmol) were combined in 5 mL DCE. Next, acetic acid (0.13 mL, 2.1 mmol) was added (apparent pH~5 when checked by wet pH paper) and the reaction was stirred at RT under argon for 1 h. At which time, NaB(O<sub>2</sub>CCH<sub>3</sub>)<sub>3</sub>H (0.286 g, 2.1 mmol) was added, and slow bubbling was observed. Reaction stirred at room temperature under argon. Once complete by TLC (EtOAc eluent) or HPLC, the reaction was diluted with CH<sub>2</sub>Cl<sub>2</sub> and transferred to a separatory funnel. The organic layer was washed with sat. NaHCO<sub>3</sub> aq. solution (2x), brine, and then organics were dried over Na<sub>2</sub>SO<sub>4</sub>. The drying agent was removed by filtration, and the organics were concentrated down to a clear oil. The residue was purified through a silica gel column, eluting with CH<sub>2</sub>Cl<sub>2</sub> to remove upper running spots (starting materials). Then, the polarity was increased with ethyl acetate from 0% to 25%, and 2-(4-chlorophenyl)-*N*-((4-methoxynaphthalen-1-yl)methyl)ethan-1-amine (**GM46-75**) was collected. Fractions were concentrated to yield 22 mg (16%) of a clear oil. HRMS (ESI-TOF, positive mode) *m/z* ([*M*+1]); Anal. Calcd. for C<sub>20</sub>H<sub>21</sub>ClNO, 326.1312; found 326.1309; RT= 3.74 min. <sup>1</sup>H NMR (300 MHz, CDCl<sub>3</sub>) δ 8.34 – 8.22 (m, 1H), 8.01 – 7.89 (m, 1H), 7.56 – 7.40 (m, 2H), 7.29 (d, *J* = 7.8 Hz, 1H), 7.25 – 7.17 (m, 2H), 7.16 – 7.06 (m, 2H), 6.71 (d, *J* = 7.8 Hz, 1H), 4.13 (s, 2H), 3.97 (s, 3H), 2.95 (t, *J* = 7.4 Hz, 2H), 2.79 (t, *J* = 7.1 Hz, 2H).

##### Method M13. Synthesis of MSU-18

In Rb flask, 4-methoxynaphthalenecarbaldehyde (0.088 g, 0.46 mmol) and 2-(4-(trifluoromethoxy)phenyl)ethan-1-amine (0.096 g, 0.46 mmol) were combined in 5 mL DCE. Next, acetic acid (0.11 mL, 1.9 mmol) was added (apparent pH~5 when checked by wet pH paper) and the reaction was stirred at RT under argon for 1 h. At which time, NaB(O<sub>2</sub>CCH<sub>3</sub>)<sub>3</sub>H (0.147 g, 0.69 mmol) was added, and slow bubbling was observed. Reaction stirred at room temperature under argon. Once complete by TLC (EtOAc eluent) or HPLC, the reaction was diluted with CH<sub>2</sub>Cl<sub>2</sub> and transferred to a separatory funnel. The organic layer was washed with sat. NaHCO<sub>3</sub> aq. solution (2x), brine, and then organics were dried over Na<sub>2</sub>SO<sub>4</sub>. The drying agent was removed by filtration, and the organics were concentrated down to a clear oil. The residue was purified through a silica gel column, eluting with $\text{CH}_2\text{Cl}_2$ to remove upper running spots (starting materials). Then, the polarity was increased with ethyl acetate from 0% to 100%, and *N*-((4-methoxynaphthalen-1-yl)methyl)-2-(4-(trifluoromethoxy)phenyl)ethan-1-amine (**MSU-18**) was collected. Fractions were concentrated to yield 87 mg (48%) of a yellow oil. HRMS (ESI-TOF, positive mode) $m/z$ ($[M+1]$); Anal. Calcd. for $\text{C}_{21}\text{H}_{21}\text{F}_3\text{NO}_2$, 376.1524; found 376.1518; RT= 4.05 min. $^1\text{H}$ NMR (400 MHz, $\text{CDCl}_3$) $\delta$ 8.26 – 8.17 (m, 1H), 7.93 – 7.84 (m, 1H), 7.40 (tt, $J$ = 6.8, 5.1 Hz, 2H), 7.22 (s, 1H), 7.18 – 7.08 (m, 2H), 7.06 – 6.99 (m, 2H), 6.64 (d, $J$ = 7.8 Hz, 1H), 4.07 (s, 2H), 3.90 (s, 3H), 2.90 (t, $J$ = 7.3 Hz, 2H), 2.75 (t, $J$ = 7.1 Hz, 2H). $^{19}\text{F}$ NMR (376 MHz, $\text{CDCl}_3$) $\delta$ -57.88 (s, 3F).

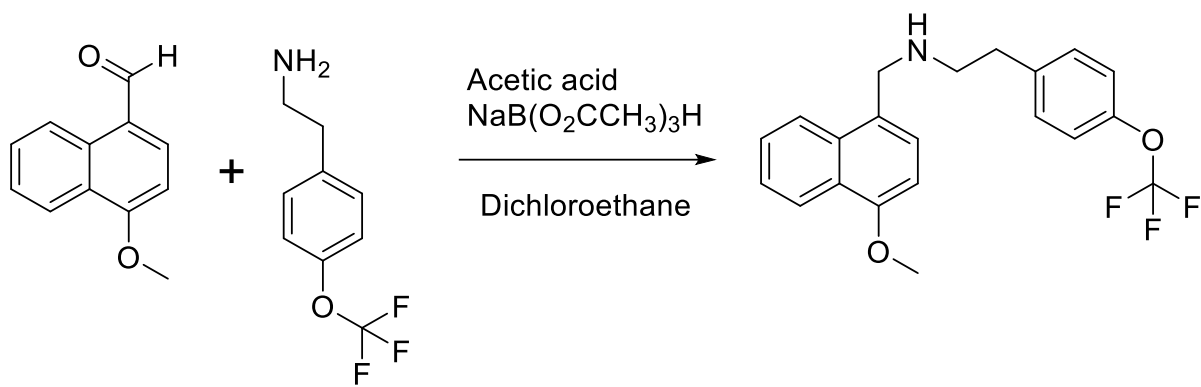

##### Method M14. Synthesis of MSU-31

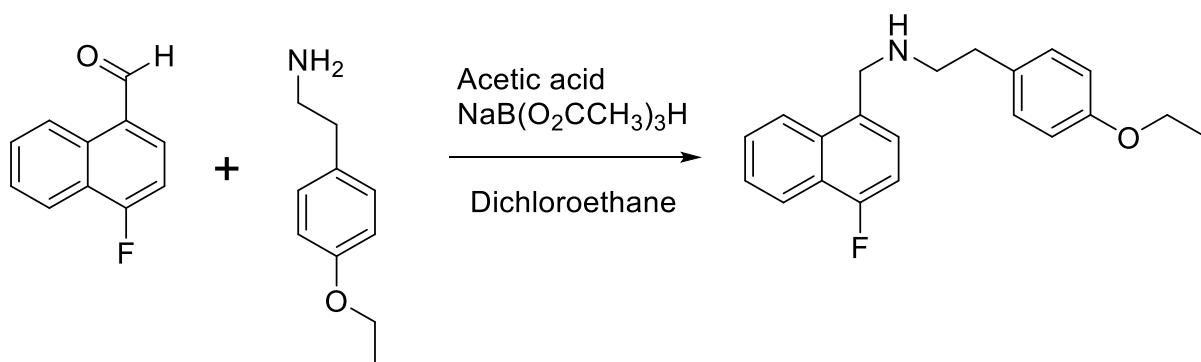

In Rb flask, 4-fluoronaphthalenecarbaldehyde (0.103 g, 0.58 mmol) and (4-ethoxyphenyl)ethanamine (0.120 g, 0.58 mmol) were combined in 5 mL DCE. Next, acetic acid (0.13 mL, 2.3 mmol) was added (apparent pH~5 when checked by wet pH paper) and the reaction was stirred at RT under argon for 1 h. At which time, $\text{NaB}(\text{O}_2\text{CCH}_3)_3\text{H}$ (0.185 g, 0.87 mmol) was added, and slow bubbling was observed. Reaction stirred at room temperature under argon. Once complete by TLC (EtOAc eluent) or HPLC, the reaction was diluted with $\text{CH}_2\text{Cl}_2$ and transferred to a separatory funnel. The organic layer was washed with sat. $\text{NaHCO}_3$ aq. solution (2x), brine, and then organics were dried over $\text{Na}_2\text{SO}_4$. The drying agent was removed by filtration, and the organics were concentrated down to a clear oil. The residue was purified through a silica gel column, eluting with $\text{CH}_2\text{Cl}_2$ to remove upper running spots (starting materials). Then, the polarity was increased with ethyl acetate from 0% to 25%, and 2-(4-ethoxyphenyl)-*N*-((4-fluoronaphthalen-1-yl)methyl)ethan-1-amine (**MSU-31**) was collected. Fractions were concentrated to yield 116 mg (58%) of a clear oil. HRMS (ESI-TOF, positive mode) $m/z$ ($[M+1]$); Anal. Calcd. for $\text{C}_{21}\text{H}_{23}\text{FNO}$, 324.1764; found 324.1760; RT= 3.64 min. $^1\text{H}$ NMR (300 MHz, MeOD) $\delta$ 8.06 (dt, $J$ = 7.8, 3.1 Hz, 1H), 8.01 – 7.89 (m, 1H), 7.60 – 7.47 (m, 2H), 7.38 (dd, $J$ = 7.9, 5.4 Hz, 1H), 7.05 (d, $J$ = 8.6 Hz, 3H), 6.77 (d, $J$ = 8.5 Hz, 2H), 4.13 (s, 2H), 3.95 (q, $J$ = 7.0 Hz, 2H), 2.88 (ddd, $J$ = 7.6, 6.6, 1.2 Hz, 2H), 2.75 (t, $J$ = 6.8 Hz, 2H), 1.34 (t, $J$ = 7.0 Hz, 3H). $^{19}\text{F}$ NMR (282 MHz, MeOD) $\delta$ -126.28 (dd, $J$ = 10.9, 5.6 Hz, 1F).

##### Method M15. Synthesis of MSU-155

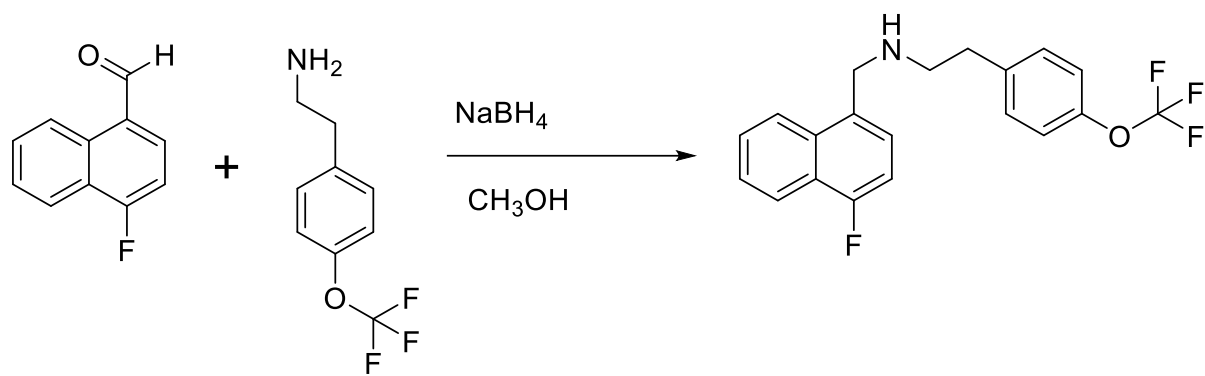

In Rb flask, 4-fluoronaphthalenecarbaldehyde (0.06 g, 0.34 mmol) and 2-(4-(trifluoromethoxy)phenyl)ethanamine (0.07 g, 0.34 mmol) were combined in 5 mL anhydrous $\text{CH}_3\text{OH}$ and the reaction was stirred at RT under argon for 12 h. At which time, $\text{NaBH}_4$ (0.019 g, 0.51 mmol) was added, and rapid bubbling was observed. Reaction stirred at room temperature under argon. Once complete by TLC (EtOAc eluent), the reaction was diluted with $\text{CH}_2\text{Cl}_2$ and transferred to a separatory funnel. The organic layer was washed with sat. $\text{NaHCO}_3$ aq. solution (2x), brine, and then organics were dried over $\text{Na}_2\text{SO}_4$. The drying agent was removed by filtration, and the organics were concentrated down to a clear oil. The residue was purified through a silica gel column, eluting with $\text{CH}_2\text{Cl}_2$ to remove upper running spots (starting materials). Then, the polarity was increased with ethyl acetate from 0% to 25%, N-((4-fluoronaphthalen-1-yl)methyl)-2-(4-(trifluoromethoxy)phenyl)ethan-1-amine (**MSU-155**) was collected. Fractions were concentrated to yield 96 mg (74%) of a clear yellow oil. HRMS (ESI-TOF, positive mode) $m/z$ ($[M+1]^+$); Anal. Calcd. for $\text{C}_{20}\text{H}_{18}\text{F}_4\text{NO}$, 364.1325; found 364.1310; RT= 3.92 min. $^1\text{H}$ NMR (400 MHz, $\text{CDCl}_3$) $\delta$ 8.10 – 8.02 (m, 1H), 7.96 (dt, $J$ = 7.3, 2.2 Hz, 1H), 7.52 – 7.41 (m, 2H), 7.27 (dd, $J$ = 7.8, 5.4 Hz, 1H), 7.17 – 7.10 (m, 2H), 7.04 (d, $J$ = 8.2 Hz, 2H), 6.99 (dd, $J$ = 10.4, 7.8 Hz, 1H), 4.12 (s, 2H), 2.92 (t, $J$ = 7.0 Hz, 2H), 2.77 (t, $J$ = 7.0 Hz, 2H). $^{19}\text{F}$ NMR (376 MHz, $\text{CDCl}_3$) $\delta$ -57.89 ($\text{OCF}_3$), -124.18 (dd, $J$ = 10.5, 5.6 Hz, 1F).

##### Method M16. Synthesis of MSU-156

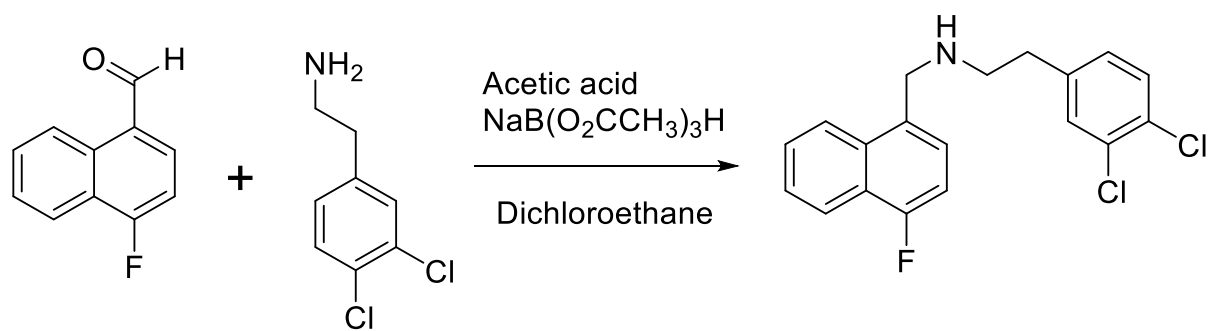

In Rb flask, 4-fluoronaphthalenecarbaldehyde (0.100 g, 0.56 mmol) and 2-(3,4-dichlorophenyl)ethanamine (0.109 g, 0.56 mmol) were combined in 5 mL anhydrous DCE. Next, acetic acid (0.13 mL, 2.25 mmol) was added (apparent pH~5 when checked by wet pH paper) and the reaction was stirred at RT under argon for 5 h. At which time, $\text{NaB}(\text{O}_2\text{CCH}_3)_3\text{H}$ (0.179 g, 0.84 mmol) was added, and rapid bubbling was observed. Reaction stirred at room temperature under argon. Once complete by TLC (EtOAc eluent) the reaction was diluted with $\text{CH}_2\text{Cl}_2$ and transferred to a separatory funnel. The organic layer was washed with sat. $\text{NaHCO}_3$ aq. solution (2x), brine, and then organics were dried over $\text{Na}_2\text{SO}_4$. The drying agent was removed by filtration, and the organics were concentrated down to a clear oil. The residue was purified through a silica gel column, eluting with $\text{CH}_2\text{Cl}_2$ to remove upper running spots (starting materials). Then, the polarity was increased with ethyl acetate from 0% to 25%, 2-(3,4-dichlorophenyl)-N-((4-fluoronaphthalen-1-yl)methyl)ethan-1-amine (**MSU-156**) was collected. Fractions were concentrated to yield 67 mg (34%) of a clear rose oil. HRMS (ESI-TOF, positive mode) $m/z$ ($[M+1]$); Anal. Calcd. for $C_{19}H_{17}Cl_2FN$, 348.0722; found 348.0724; RT= 3.80 min. $^1H$ NMR (400 MHz, $CDCl_3$) $\delta$ 8.06 – 7.97 (m, 1H), 7.92 (dq, $J$ = 7.3, 2.7 Hz, 1H), 7.50 – 7.39 (m, 2H), 7.25 – 7.11 (m, 2H), 6.95 (dd, $J$ = 10.3, 7.8 Hz, 1H), 6.89 (dd, $J$ = 8.2, 2.1 Hz, 1H), 4.06 (s, 2H), 2.85 (t, $J$ = 7.0 Hz, 2H), 2.66 (t, $J$ = 7.0 Hz, 2H). $^{19}F$ NMR (376 MHz, $CDCl_3$) $\delta$ -109.90 – -131.06 (m, 1F).

**LC-MS method.** Liquid Chromatography- Mass Spectrometry was performed on an Agilent 1290 infinity coupled to Agilent 6538 Ultra High-Definition Quadrupole Time of Flight (UHD-QToF) instrument. Separations were achieved by using reverse phase Waters Acquity UPLC HSS T3 1.8μm (2.1 X 100mm) column from Waters (Milford, USA). The LCMS Optima grade solvents were purchased from Fischer Scientific. Water containing 0.1% formic acid was used as mobile phase A and acetonitrile containing 0.1% formic acid was used as mobile phase B. The injection volume was set at 1 μL. Samples were injected in a gradient of 95% mobile phase A and 5% mobile phase B in the initial condition to 5% mobile phase A and 95% mobile phase B in 9 min. The ratio was held at that composition for an additional 3 min and switched back to the initial condition at 12 min. The MS data acquisition was performed from 50-1000$m/z$ at a 1.0 spectra/sec scan rate. The source gas temperature was set at 350 $^{\circ}C$ with a flow of 8 L/min. The nebulizer gas was set at 55 psig. The capillary voltage was set at 3500 volts with fragmentor at 100, skimmer at 45 and octopole RF 500 volts. Prior to sample runs, the instrument was calibrated using Agilent low mass calibrant solution.

### SUPPLEMENTARY FIGURES

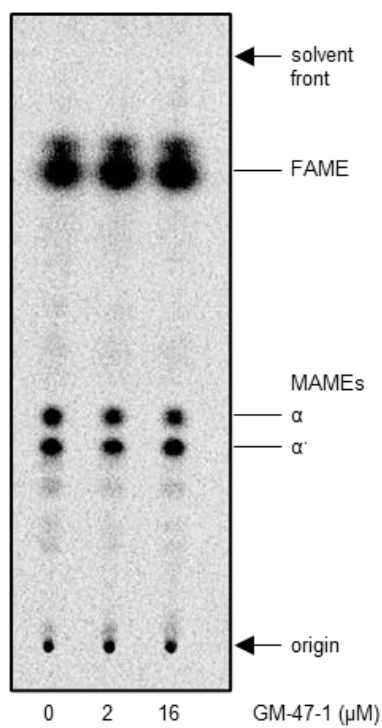

**Figure S1. GM-47-1 does not affect *de novo* mycolic acid biosynthesis.** TLC analysis of [$^{14}\text{C}$]-labeled methyl esters of whole cell acyl chains. The same amount of radioactivity was loaded for each sample. The TLC was visualized by phosphor imaging. Each TLC is representative of at least two biological replicates. FAME, fatty acid methyl ester; MAME, mycolic acid methyl ester.

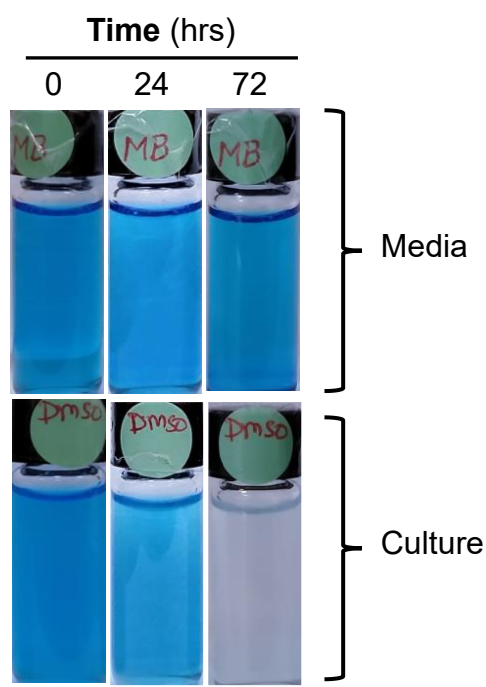

**Figure S2. Oxygen consumption decolorizes methylene blue.** Exponentially grown culture was resuspended at a density of 0.3 measured at 600 nm. Glass vials after adding the culture were and methylene blue at 0.001% were tightly sealed and incubated at 32°C in anaerobic jar. Upper lane contain only media (no cells) as was used as colour control, lower lane contained culture.

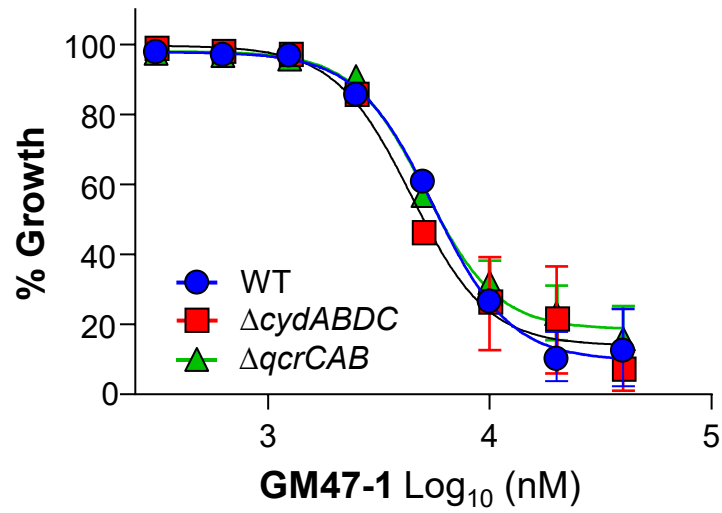

**Figure S3. Growth inhibitory potency of GM47-1 remains unaltered by loss of either respiratory terminal oxidase.** MIC of GM47-1 was determined using *M. abscessus* ATCC 19977 (wild type) strain and two knockout strains representing loss of either terminal oxidase i.e. *cydABDC* and *qcrCAB* operon.

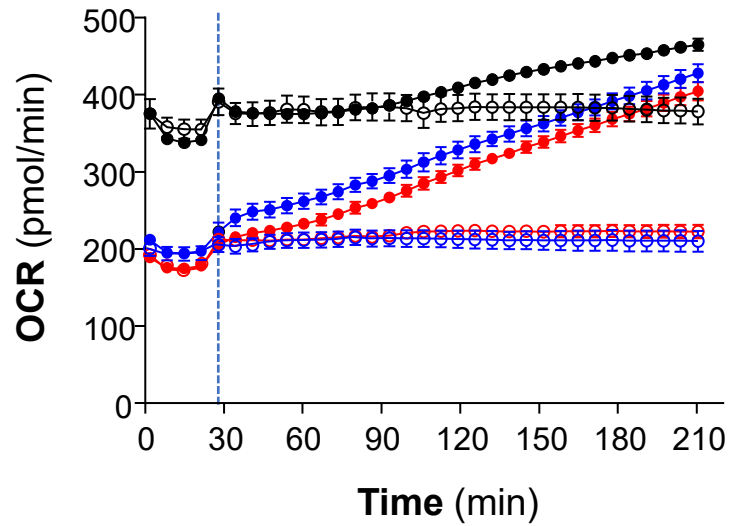

**Figure S4. Respiratory acceleration is mediated by cytochrome *bcc:aa<sub>3</sub>*.** Effect of GM47-1 on comparison of OCR using parental strain (blue), $\Delta cydABDC$ (red) and $\Delta qcrCAB$ (black). GM47-1 was injected by 2× for GM47-1, as described by filled circles. Raw data points are shown for comparison and injection timepoint is shown by dotted line.

### SUPPLEMENTARY TABLES

**Table S1. ATP depleting hits obtained from LOPAC<sup>®1280</sup> hits are nonspecific towards *cyt-bcc:aa<sub>3</sub>*.** Absolute ATP depletion is defined as ratio of ATP value to basal ATP levels. Bedaquiline was used as a control drug. ATP dose response of LOPAC<sup>®1280</sup> hits were performed against *M. abscessus* Δ*cydABDC* and parental strain. Bedaquiline was used as a control drug. ATP IC<sub>50</sub> is defined as effective concentration required for half the maximal decrease in bioluminescence (ATP).

| Drug Name | Target | Absolute ATP depletion (Log<sub>2</sub>-fold decrease) | ATP IC<sub>50</sub> (μM) against *M. abscessus* |  |
| --- | --- | --- | --- | --- |
|  |  |  | Δ*cydABDC* | Wild type |
| Ofloxacin | DNA gyrase (antibiotic) | 1.18 | >40 | >40 |
| Lomefloxacin HCl | DNA gyrase (antibiotic) | 1.08 | >40 | >40 |
| Calcimycin | Intracellular calcium | 2.22 | 2.5 | 3.0 |
| AC-93253 iodide | Hormone agonist | 1.24 | 19.2 | 19.4 |
| 5-Fluorouracil | Thymidylate synthetase inhibitor (cell cycle) | 1.55 | 25.9 | 20.9 |
| Mitoxantrone dihydrochloride | DNA metabolism inhibitor | 2.06 | 10.5 | 7.1 |
| Ruthenium red | Ion pump inhibitor (mitochondrial uniporter) | 1.10 | 12.2 | 10.0 |
| Control drug (BDQ) | ATP synthase inhibitor | 1.85 | 0.04 | 0.05 |

**Table S2. Validation of hits from LOPAC<sup>®1280</sup> library screen for ATP perturbations against Mabs wild type.** All 9 hits (drugs) were identified to triggers ATP increase relative to basal levels against Mabs wild type. One among nine hits was able to inhibit growth of *M. abscessus*. BDQ, a F<sub>1</sub>F<sub>0</sub> ATP synthase inhibitor was used as a control drug which inhibits ATP synthesis and growth in *M. abscessus*. Basal levels: ATP basal levels in untreated controls. EC<sub>50</sub> is defined as effective concentration required for half the maximal increase in bioluminescence (ATP).

| Drug name | Target | ATP<br>(× fold<br>increase) | ATP<br>EC<sub>50</sub> (μM) | MIC<sub>50</sub><br>(μM) |
| --- | --- | --- | --- | --- |
| Cephalexin | Peptidoglycan<br>(antibiotic) | 2.2 | 1.8 | >40 |
| Cefaclor |  | 3.5 | 1.8 | >40 |
| Ceftriaxone |  | 1.8 | 1.5 | >40 |
| Cefotaxime |  | 2.6 | 13.1 | >40 |
| Cephadrine |  | 3.1 | 1.1 | >40 |
| Vancomycin |  | 3.8 | 8.8 | >40 |
| SBI-0087702 | Mitochondrial ATF<br>translocator | 52.7 | 10.1 | 5.6 |
| Demeclocycline | Ribosome<br>(antibiotic) | 1.6 | 3.2 | >40 |
| NS8593 | Ca<sup>2+</sup>-activated K<sup>+</sup><br>channels | 4.8 | 23.1 | >40 |

**Table S3. Growth inhibitory potency of GM47-1 against Mabs clinical isolates.** The MIC<sub>50</sub> values indicate the minimum drug concentration at which 50% of bacterial growth was inhibited in a full dose-response curve.

| Strain | rrl mutation<br>(2058/2059) | erm(41)<br>sequevar | Morph-<br>otype | GM47-1 | BDQ | Antibiotic<br>resistance<br>profile |
| --- | --- | --- | --- | --- | --- | --- |
|  |  |  |  | MIC<sub>50</sub> (μM) |  |  |
| WT<br>(CIP104536) | - | - | - | 4.7 | 0.13 | AMK<sup>S</sup>,<br>CLR<sup>S</sup>, LZD<sup>S</sup> |
| LMS | Wt | T28 | Rough | 7.2 | 0.54 | AMK<sup>S</sup>,<br>CLR<sup>R</sup>, LZD<sup>S</sup> |
| HHJ | Wt | T28 | Smooth | 4.8 | 0.21 | AMK<sup>S</sup>,<br>CLR<sup>S</sup>, LZD<sup>R</sup> |
| HNS | Wt | T28 | Rough | 3.0 | 0.07 | AMK<sup>S</sup>,<br>CLR<sup>R</sup>, LZD<sup>R</sup> |
| KDS | Wt | T28 | Smooth | 2.2 | 0.04 | AMK<sup>S</sup>,<br>CLR<sup>S</sup>, LZD<sup>R</sup> |
| SKS | Wt | C28 | Smooth | 2.4 | 0.10 | ND |
| KBS | Wt | C28 | Rough | 1.0 | 0.04 | ND |
| KMR | C/A | T28 | Rough | 3.0 | 0.10 | AMK<sup>R</sup>,<br>CLR<sup>R</sup>, LZD<sup>S</sup> |
| KYS | C/A | T28 | Rough | 6.2 | 0.06 | ND |
| KHA | C/A | C28 | Smooth | 4.6 | 0.03 | ND |
| LMK | G/A | C28 | Smooth | 2.4 | 0.50 | ND |
| LJS | Wt | T28 | Rough | 4.5 | 0.12 | ND |

BDQ: bedaquiline; AMK: amikacin; CLR: clarithromycin; LZD: linezolid; R: resistant; S: susceptible; ND: not determined

**Table S4. Cytotoxicity of GM47-1 derivatives against HepG2 cells.** To determine cytotoxicity, HepG2 cells were treated with two-fold serially diluted dose range of compounds or tamoxifen. Well with untreated cells containing vehicle (1% DMSO) and tamoxifen were used as negative and positive controls respectively.

| **Compound code** | **MIC<sub>50</sub>** | **CC<sub>50</sub>** | **SI (CC<sub>50</sub>/MIC<sub>50</sub>)** |
| --- | --- | --- | --- |
| GM47-1 | 3.9 | 31.5 | 8.07 |
| MSU-155 | 0.21 | >50 | >238 |
| MSU-156 | 0.18 | 47.4 | 263.3 |
| Tamoxifen | - | 14.9 | - |
